# DeCompress: tissue compartment deconvolution of targeted mRNA expression panels using compressed sensing

**DOI:** 10.1101/2020.08.14.250902

**Authors:** Arjun Bhattacharya, Alina M. Hamilton, Melissa A. Troester, Michael I. Love

## Abstract

Targeted mRNA expression panels, measuring up to 800 genes, are used in academic and clinical settings due to low cost and high sensitivity for archived samples. Most samples assayed on targeted panels originate from bulk tissue comprised of many cell types, and cell-type heterogeneity confounds biological signals. Reference-free methods are used when cell-type-specific expression references are unavailable, but limited feature spaces render implementation challenging in targeted panels. Here, we present *DeCompress*, a semi-reference-free deconvolution method for targeted panels. *DeCompress* leverages a reference RNA-seq or microarray dataset from similar tissue to expand the feature space of targeted panels using compressed sensing. Ensemble reference-free deconvolution is performed on this artificially expanded dataset to estimate cell-type proportions and gene signatures. In simulated mixtures, four public cell line mixtures, and a targeted panel (1199 samples; 406 genes) from the Carolina Breast Cancer Study, *DeCompress* recapitulates cell-type proportions with less error than reference-free methods and finds biologically relevant compartments. We integrate compartment estimates into *cis*-eQTL mapping in breast cancer, identifying a tumor-specific *cis*-eQTL for *CCR3* (C-C Motif Chemokine Receptor 3) at a risk locus. *DeCompress* improves upon reference-free methods without requiring expression profiles from pure cell populations, with applications in genomic analyses and clinical settings.

## INTRODUCTION

Academic and clinical settings have prioritized the collection of tissue samples of mixed cell types for molecular profiling and biomarker studies (1–3). Bulk tissue, especially from cancerous tumors, is comprised of different cell types, many rare, and each contributing varied biological signal to an assay (e.g. mRNA expression) (4, 5). This cell-type heterogeneity makes it difficult to distinguish variability that reflects shifts in cell populations from variability that reflects changes in cell-type-specific expression (6). Since RNA-seq technology was developed, cell-type deconvolution from mRNA expression has become important in genetic and genomic association studies: either using compositions in regression models as covariates to adjust for the association between cell type and phenotype (7–10), or using them as inputs to solve for cell-type specific quantities (11, 12). Cell-type deconvolution methods can be reference-based (supervised) (13–19) or reference-free (unsupervised) (20–26), depending on whether cell-type-specific expression profiles are available for the component cell-types. When reference panels are unavailable, as in understudied tissues or populations (27), reference-free deconvolution is the only viable option. Even in cases where reference expression profiles are available, reference-based methods may provide inaccurate proportion estimates if the mixed tissue and references represent different clinical settings or phenotypes (28).

Given the advent of single-cell technologies and studies into cell trajectories, the concept of cell types in bulk tissue has been debated (29). Especially in perturbed or diseased tissues, like cancer, individual cells may present in different states, or various cells of possibly different identities may contribute, in aggregate, to the same biological process and have similar molecular profiles (30–32). While previous reference-free methods rely on searching the feature space for compartment-specific molecular features from the entire transcriptome and thus require a large feature space (22, 24–26), reference-free deconvolution methods can, with fewer assumptions, identify tissue compartments, or isolated units of a tissue that represent either a biological process or a cell type (33). Thus, reference-free methods have important advantages over reference-based methods but may require a large number of features for optimal performance (25, 34).

Many important datasets may have fewer expression targets than those required for existing reference-free deconvolution methods. Targeted mRNA expression assays are optimized for gene expression quantification in samples stored clinically and use a panel of up to 800 genes without requiring cDNA synthesis or amplification steps (35–37). These technologies offer key advantages in sensitivity, technical reproducibility, and strong robustness for profiling formalin-fixed, paraffin-embedded (FFPE) samples (35, 38). Given these advantages, targeted expression profiling is increasingly being used for molecular studies (36, 37, 39–42), especially prospective studies involving FFPE samples stored over several years (43) and diagnostic assays in clinical settings (3, 44). Due to its viability in diagnostics, it is important to identify reference-free deconvolution methods that overcome the need for searching for compartment-specific genes from the assay’s feature space (22, 24–26), given the limited feature space in targeted panels.

Previous groups have proposed methods for efficiently reconstructing full gene expression profiles from sparse measurements of the transcriptome, borrowing techniques from image reconstruction using compressed sensing (45, 46) and machine learning (47–50). For example, Cleary *et al* developed a blind compressed sensing method that recovers gene expression from multiple composite measurements of the transcriptome (up to 100 times fewer measurements than genes) by using modules of interrelated genes in an unsupervised manner. Another imputation method by Viñas *et al* (51) used recent machine learning methodology (52) to provide efficient and accurate transcriptomic reconstruction in healthy, unperturbed tissue from the Genotype-Tissue Expression (GTEx) Project (53, 54). The performance of these methods provides a promising avenue to expand the feature space of targeted panels, rendering them more applicable for reference-free deconvolution methods.

Here, we present *DeCompress*, a semi-reference-free deconvolution method for targeted panels. *DeCompress* requires a reference RNA-seq or microarray dataset from the same bulk tissue assayed by the targeted expression panel to train a compressed sensing model to expand the feature space in a targeted panel. We show the advantages of using *DeCompress* over other reference-free methods with simulation analyses and real data applications. Lastly, we examine the impact of tissue compartment deconvolution on downstream analyses, such as *cis*-eQTL analysis using expression data from the Carolina Breast Cancer Study (CBCS) (55). *DeCompress* is available freely as an R package on GitHub at https://github.com/bhattacharya-a-bt/DeCompress.

## MATERIAL AND METHODS

### The Decompress algorithm

*DeCompress* takes in two expression matrices from similar bulk tissue as inputs: the *target* expression matrix from a targeted panel of gene expression with *n* samples and *k* genes, and a *reference* expression matrix from an RNA-seq and microarray panel with *N* samples and *K > k* genes. Ideally, both the target and reference expression matrices should be on the raw expression scale (not log-transformed), as we presume the total RNA abundance for a given gene in bulk tissue is a linear combination of that gene’s compartment-specific RNA abundance. We refer to DeCompress as a semi-reference-free method, as it requires a reference expression matrix but not compartment-specific expression profiles (as in reference-based methods). For a user-defined number of compartments, *DeCompress* outputs compartment proportions for all samples in the target and the compartment-specific expression profiles for the genes used in deconvolution. The method follows three general steps, as detailed in **Figure 1**: (1) selection of the compartment-specific genes from the reference, (2) compressed sensing to expand the targeted panel to a *DeCompressed* expression matrix with these compartment-specific genes, and (3) ensemble deconvolution on the DeCompressed dataset. Full mathematical and algorithmic details for *DeCompress* are provided in **Supplemental Methods**. *DeCompress* is available as an R package on GitHub (https://github.com/bhattacharya-a-bt/DeCompress).

**Figure 1:**
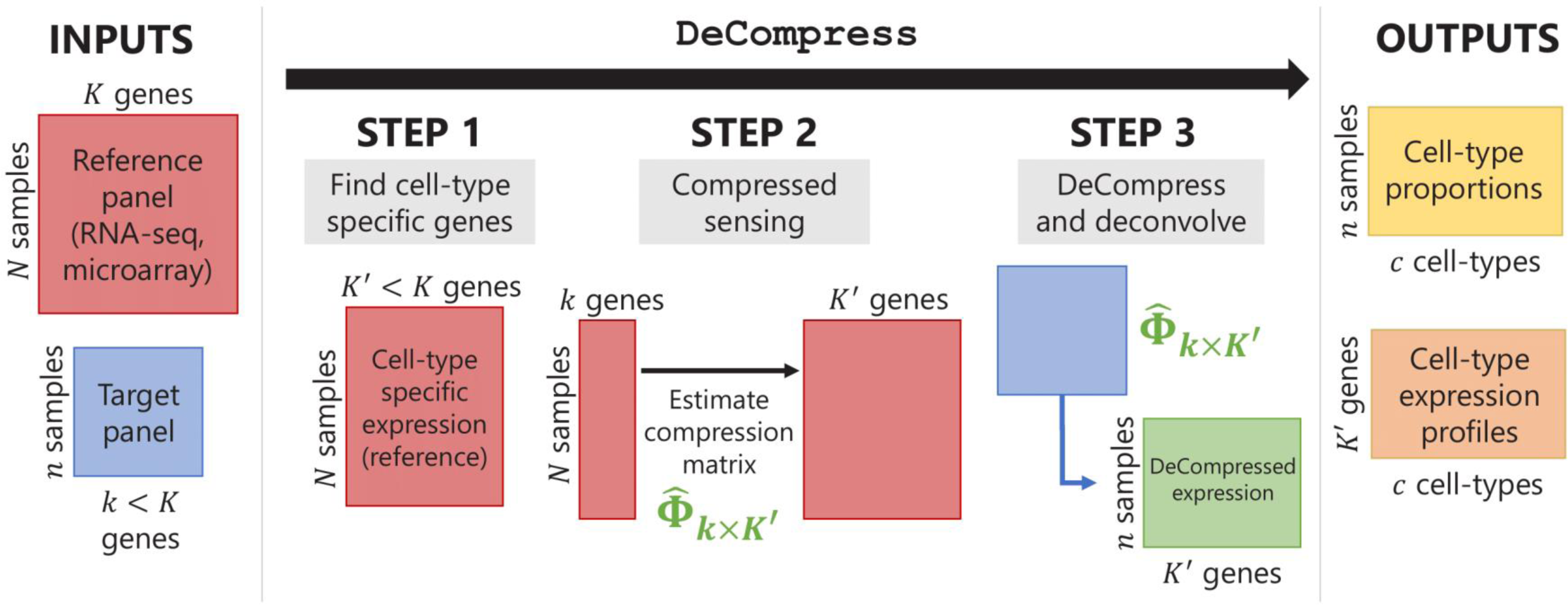
Schematic for the DeCompress algorithm. *DeCompress* takes in a *reference* RNA-seq or microarray matrix with *N* samples and *K* genes, and the *target* expression with *n* samples and *k < K* genes. The algorithm has three general steps: (1) finding the *K*′ *< K* genes in the reference that are cell-type specific, (2) training the compressed sensing model that projects the feature space in the target from *k* genes to the *K′* cell-type specific genes, and (3) decompressing the target to an expanded dataset and deconvolving this expanded dataset. *DeCompress* outputs cell-type proportions and cell-type specific profiles for the *K′* genes.

The first step of *DeCompress* is to use the reference dataset to find a set of *K*′ *< K* genes that are representative of different compartments that comprise the bulk tissue. These *K′* genes, called the compartment-specific genes, can be supplied by the user if prior gene signatures can be applied. If any such gene signatures are not available, *DeCompress* borrows from previous reference-free methods to determine this set of genes (*Linseed* (22) or *TOAST* (25)). If the user cannot determine the total number of compartments, using the reference, the number of compartments can be estimated by assessing the cumulative total variance explained by successive singular value decomposition modes.

After a set of compartment-specific genes are determined, *DeCompress* uses the reference to infer a model that predicts the expression of each of these compartment-specific genes from the genes in the target. Predictive modeling procedures borrow ideas from compressed sensing (45, 46, 56), a technique that was developed to reconstruct a full image from sparse measurements of it: the estimation procedure can be broken down into solving a system of equations using either linear or non-linear regularized optimization, with options for parallelization when the sample size of the reference dataset is large. These optimization methods are detailed in **Supplemental Methods**. The predictive models are curated into a *compression* matrix, which is then used to expand the original target (with *k < K*′ *< K* genes) into the artificially *DeCompressed* expression matrix (with the *K’* compartment-specific genes). In practice, we observed that regularized linear regression (lasso, ridge, or elastic net regression (48)) provides the best prediction of gene expression (**Supplemental Figure S1**), and the user may either model the gene expression using the traditional Gaussian family or assume that the errors follow a Poisson distribution to account for the scale of the original data (not log-transformed).

Lastly, ensemble deconvolution is performed on the DeCompressed expression matrix to estimate (1) compartment proportions on the samples in the target, and (2) the compartment-specific expression profiles for the *K*′ genes used in deconvolution. Several options for reference-free deconvolution are provided in *DeCompress*. We also provide options that uses a reference-based method, *unmix* from the DESeq2 package (57), based on compartment expression profiles estimated from the reference RNA-seq or microarray dataset (i.e. an approximate compartment expression profile is estimated from a non-negative matrix factorization of the reference dataset). Estimates from the method that best recovers the DeCompressed expression matrix is chosen. **Supplemental Table S1** provides summaries of the methods employed in *DeCompress*.

### Benchmarking analysis

Using simulations and published datasets, we benchmarked *DeCompress* against five other reference-free methods: *deconf* (20), *CellDistinguisher* (26), *Linseed* (22), *DeconICA* (24), and iterative non-negative matrix factorization with feature selection using TOAST (25) (see **Supplemental Table S1**). All these datasets provide a matrix of known compartment proportions. To measure the performance of each method, we calculate the error between the estimated and true compartment proportions as the mean square error (MSE) (i.e. the mean row-wise MSE between the two matrices). We also permute the columns the estimated matrix (corresponding to compartments) to align compartments accordingly between the known and estimated proportions to minimize the MSE for each method.

#### In-silico mixing with GTEx

We performed *in-silico* mixing experiments using expression data from the Genotype-Tissue Expression (GTEx) Project (dbGAP accession number phs000424.v7.p2) (53, 54). Here, we obtained median transcripts per kilobase million (TPM) data for four tissue types: mammary tissue, EBV-transformed lymphocytes, transformed fibroblasts, and subcutaneous adipose. We randomly generated compartment proportions for each of these tissue types and simulated mixed RNA-seq expression data for 200 samples. We then scaled these mixed expression profiles with multiplicative noise randomly generated from a Normal distribution with 0 mean and standard deviations of 4 and 8. We then generated 25 pseudo-targeted expression panels by randomly selecting 200, 500, and 800 of the genes with mean and standard deviations above the median mean and standard deviations of all genes. For benchmarking, we randomly select 100 samples for the target matrix. For *DeCompress*, the simulated RNA-seq data on the other 100 samples are used as the reference matrix. We added more normally-distributed multiplicative noise with zero mean and unit variance to simulate a batch difference between the reference and target matrix. For comparison to compartments with dissimilar expression profiles, we repeated these simulations for four other tissues: mammary tissue, pancreas, pituitary, and whole blood. Full details for this simulation framework are provided in **Supplemental Methods**.

#### Existing mixing experiments

We also benchmarked *DeCompress* in four published mixing experiments: (1) microarray expression for mixed rat brain, liver, and lung biospecimens (GEO Accession Number: GSE19830), commonly used as a benchmarking dataset in deconvolution studies (*N* = 42) (11), (2) RNA-seq expression (GSE123604) for a mixture of breast cancer cells, fibroblasts, normal mammary cells, and Burkitt’s lymphoma cells (*N* = 40) (23), (3) microarray expression (GSE97284) for laser capture micro-dissected prostate tumors (*N* = 30) (58), and (4) RNA-seq expression (GSE64098) for a mixture of two lung adenocarcinoma cell lines (*N* = 40) (59, 60). As in the in-silico mixing using GTEx data, we generated pseudo-targeted panels by randomly selecting 200, 500, and 800 of the genes with mean and standard deviations above the median mean and standard deviations of all genes. For the rat mixture dataset, we used 30 of the 42 samples as a reference microarray matrix (with multiplicative noise, as in GTEx) and deconvolved on the remaining 12 samples in the target matrix. In the remaining three datasets, we obtained normalized RNA-seq reference matrices from The Cancer Genome Atlas: TCGA-BRCA breast tumor expression for the breast cancer cell line mixture, TCGA-PRAD prostate tumor expression for the prostate tumor microarray study, and TCGA-LUAD for the lung adenocarcinoma mixing study. These datasets are summarized in **Supplemental Table S2**.

### Applications in Carolina Breast Cancer Study (CBCS) data

We lastly used expression data from the Carolina Breast Cancer Study for validation and analysis (55). Paraffin-embedded tumor blocks were requested from participating pathology laboratories for each samples, reviewed, and assayed for gene expression using the NanoString nCounter system, as discussed previously (43). As described before (10, 61), the expression data (406 genes and 11 housekeeping genes) was pre-processed and normalized using quality control steps from the *NanoStringQCPro* package, upper quartile normalization using *DESeq2* (57, 62), and estimation and removal of unwanted technical variation using the *RUVSeq* and *limma* packages (63, 64). The resulting normalized dataset comprised of samples from 1,199 patients, comprising of 628 women of African descent (AA) and 571 women of European descent (EA). A study pathologist analyzed tumor microarrays (TMAs) from 148 of the 1,199 patients to estimate area of dissections originating from epithelial tumor, intratumoral stroma, immune infiltrate, and adipose tissue (10). These compartment proportions of the 148 samples were used for benchmarking of *DeCompress* against other reference-free methods.

Date of death and cause of death were identified by linkage to the National Death Index. All diagnosed with breast cancer have been followed for vital status from diagnosis until date of death or date of last contact. Breast cancer-related deaths were classified as those that listed breast cancer (International Statistical Classification of Disease codes 174.9 and C-50.9) as the underlying cause of death on the death certificate. Of the 1,199 samples deconvolved, 1,153 had associated survival data with 330 total deaths, 201 attributed to breast cancer.

#### Over-representation and gene set enrichment analysis

We conducted over-representation (ORA) and gene set enrichment analysis (GSEA) to identify significantly enriched gene ontologies using *WebGestaltR* (65). Specifically, we considered biological process ontologies categorized by The Gene Ontology Consortium (66, 67) at FDR-adjusted *P* < 0.05.

#### Survival analysis

Here, we defined a relevant event as a death due to breast cancer. We aggregated all deaths not due to breast cancer as a competing risk. Any subjects lost to follow-up were treated as right-censored observations. We built cause-specific Cox models (68) by modeling the hazard function of breast cancer-specific mortality with the following covariates: race, PAM50 molecular subtype (69), age, compartment-specific proportions, and an interaction term between molecular subtype and compartment proportion. We compared these compartment-specific survival models with the nested baseline model that did not include compartment proportions using partial likelihood ratio tests. We tested for the statistical significance of parameter estimates using Wald-type tests, adjusting for multiple testing burden using the Benjamini-Hochberg procedure at a 10% false discovery rate (70).

#### eQTL analysis

CBCS genotype data is measured on the OncoArray. Approximately 50% of the SNPs for the OncoArray were selected as a “GWAS backbone” (Illumina HumanCore), which aimed to provide high coverage for many common variants through imputation. The remaining SNPs were selected from lists supplied by six disease-based consortia, together with a seventh list of SNPs of interest to multiple disease-focused groups. Approximately 72,000 SNPs were selected specifically for their relevance to breast cancer. The sources for the SNPs included in this backbone, as well as backbone manufacturing, calling, and quality control, are discussed in depth by the OncoArray Consortium (71, 72). All samples were imputed using the October 2014 (v.3) release of the 1000 Genomes Project (73) as a reference panel in the standard two-stage imputation approach, using *SHAPEIT2* for phasing and *IMPUTEv2* for imputation (74–76). All genotyping, genotype calling, quality control, and imputation was done at the DCEG Cancer Genomics Research Laboratory (71, 72).

From the provided genotype data, we excluded variants (1) with a minor frequency less than 1% based on genotype dosage and (2) that deviated significantly from Hardy-Weinberg equilibrium at *P* < 10^−8^ using the appropriate functions in *PLINK v1.90b3* (77). Finally, we intersected genotyping panels for the AA and EA samples, resulting in 5,989,134 autosomal variants. We excluded 334,391 variants on the X chromosome. CBCS genotype data was coded as dosages, with reference and alternative allele coding as in the National Center for Biotechnology Information’s Single Nucleotide Polymorphism Database (dbSNP) (78).

As previously described (10), using the 1,199 samples (621 AA, 578 EA) with expression data, we assessed the additive relationship between the gene expression values and genotypes with linear regression analysis using *MatrixeQTL* (79). We consider a baseline linear model with log-transformed gene expression of a gene of interest as the dependent variable, SNP dosage as the primary predictor of interest, and the following covariates: age, BMI, post-menopausal status, and the first 5 principal components of the joint AA and EA genotype matrix. We also considered a compartment-specific interaction model that adds compartment proportion from *DeCompress* and an interaction term between the SNP dosage and compartment proportion (8, 9). This interaction model subtly changes the interpretation of the main SNP dosage effect, representing an estimate of the eQTL effect size at 0% compartment-specific cells. Thus, we recover compartment-specific eQTLs by testing the interaction effect, which measures how the magnitude of an eQTL differs between the two cell types. The interaction model was fit using *MatrixeQTL*’s linear-cross implementation. It is important to note that we model the log-transformed expression here, as existing methods for modeling expression on genotype do not support interaction terms (80–82).

We compared eQTLs mapped in CBCS here with eQTLs in GTEx. We downloaded healthy tissue eQTLs from the Genotype-Tissue Expression (GTEx) Project and cross-referenced eGenes and corresponding eSNPs between CBCS and GTEx in healthy breast mammary tissue, EBV-transformed lymphocytes, transformed fibroblasts, and subcutaneous adipose tissue. We considered these tissues mainly due to their high relative composition in bulk breast tumor samples, as shown previously in many studies (23, 83–85). The Genotype-Tissue Expression (GTEx) Project was supported by the Common Fund of the Office of the Director of the National Institutes of Health, and by NCI, NHGRI, NHLBI, NIDA, NIMH, and NINDS. The data used for the analyses described in this manuscript were obtained from the GTEx Portal on 05/14/20. We also downloaded iCOGs GWAS summary statistics for breast cancer risk (86–88) to assess any overlap between CBCS eQTLs and GWAS-detected risk variants.

## RESULTS

### Overview of the DeCompress algorithm

*DeCompress* takes in two expression matrices from similar bulk tissue as inputs: an expression matrix from a targeted panel of gene expression with *n* samples and *k* genes, and an expression matrix from an RNA-seq and microarray panel with *N* samples and *K > k* genes. For shorthand, we will refer to RNA-seq or microarray panel as the *reference* and the targeted expression panel as the *target*. *DeCompress* outputs tissue compartment proportions for a user-defined number of all samples in the target and the compartment-specific expression profiles for the genes used in deconvolution. The method follows three general steps, as detailed in **Figure 1**: (1) feature selection of the compartment-specific genes from the reference, (2) compressed sensing to expand the targeted panel to a *DeCompressed* expression matrix with these compartment-specific genes, and (3) ensemble deconvolution on the DeCompressed dataset using existing reference-free methods. We provide further details about *DeCompress* in **Methods** and full mathematical and algorithmic details in **Supplemental Methods**.

### Benchmarking DeCompress against other reference-free deconvolution methods

We benchmarked *DeCompress* performance across 6 datasets (see **Supplemental Table S2**): (1) *in-silico* mixing experiments using tissue-specific expression profiles from the Genotype-Tissue Expression (GTEx) Project (53, 54), (2) expression from 4 published datasets with known compartment proportions (11, 23, 58, 59), and (3) and tumor expression from the Carolina Breast Cancer Study (43, 55). We compared the performance of *DeCompress* against 5 other reference-free deconvolution methods (summarized in **Supplemental Table S1**): *deconf* (20), *Linseed* (22), *DeconICA* (24), iterative non-negative matrix factorization with feature selection using *TOAST* (*TOAST + NMF*) (25), and *CellDistinguisher* (26). Estimated compartment proportions are compared to simulated or reported true compartment proportions with the mean square error (MSE) between the two matrices (see **Methods**). In total, we observed that *DeCompress* recapitulates compartment proportions with the least error compared to reference-free deconvolution methods.

#### In-silico GTEx mixing

We generated artificial targeted panels by mixing median tissue specific expression profiles from GTEx *in-silico* with randomly simulated compartment proportions for mammary tissue, EBV-transformed lymphocytes, transformed fibroblasts, and subcutaneous adipose. We added multiplicative noise to the mixed expression to simulate measurement error and contributions to the bulk expression signal from other sources (see **Methods**). **Figure 2A** shows the performance of *DeCompress* compared to other reference-free methods across 25 simulated targeted panels of increasing numbers of genes on the simulated targeted panels. In general, we find that *DeCompress* gives more accurate estimates of compartment proportions than the other 5 methods at both settings for multiplicative noise. As the number of genes in the targeted panel increased, the difference in MSE between *DeCompress* and the other methods remains largely constant. *Linseed* and *DeconICA*, methods that search for mutually independent axes of variation that correspond to compartments, consistently perform poorly on these simulated datasets, possibly due to the relative similarity between the expression profiles for these compartments and the small number of genes on the targeted panels. *deconf*, *TOAST + NMF* (matrix factorization-based methods) and *CellDistinguisher* (topic modeling) perform similarly to one another and only moderately worse in comparison to *DeCompress*.

**Figure 2:**
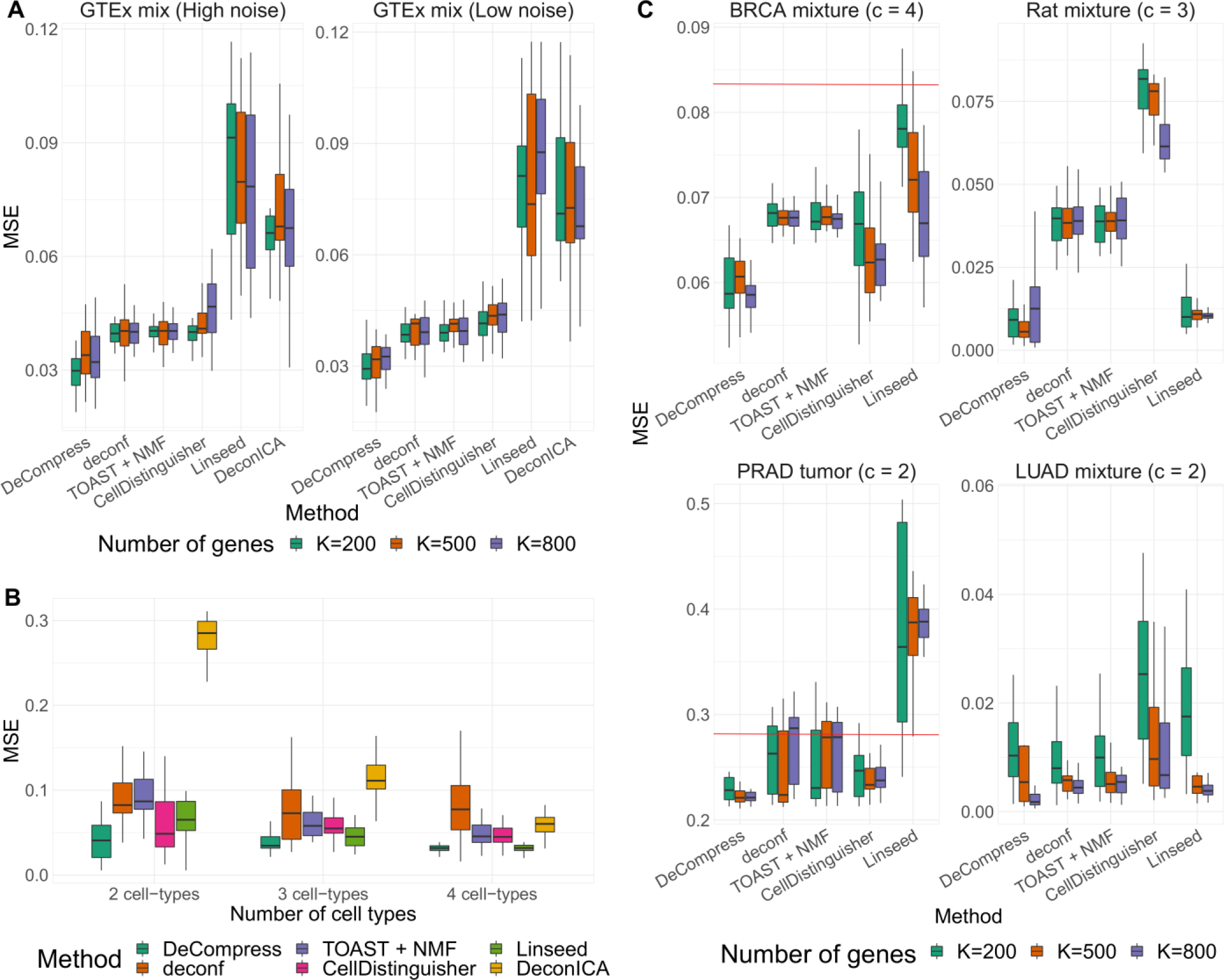
Benchmarking results for in-silico GTEx mixing experiments and real data examples. **(A)** Boxplots of mean square error (*Y*-axis) between true and estimated cell-type proportions in *in-silico* GTEx mixing experiments across various methods (*X*-axis), with 25 simulated datasets per number of genes. GTEx mixing was done at two levels of multiplicative noise, such that errors were drawn from a Normal distribution with zero mean and standard deviation 8 (left) and 4 (right). Boxplots are colored by the number of genes in each simulated dataset. **(B)** Boxplots of MSE (*Y*-axis) between true and estimated cell-type proportions over 25 simulated GTEx mixed expression datasets with 500 genes, multiplicative noise drawn from a Normal distribution with zero mean and standard deviation 10, and 2 (left), 3 (middle), and 4 (right) different cell-types. Boxplots are collected by the reference-free method tested. **(C)** Boxplots of mean square error (*Y*-axis) between true and estimated cell-type proportions in 25 simulated targeted panels of 200, 500, 800, and 1,000 genes (*X*-axis), using four different datasets: breast cancer cell-line mixture (top-left) (23), rat brain, lung, and liver cell-line mixture (top-right) (11), prostate tumor samples (bottom-left) (58), and lung adenocarcinoma cell-line mixture (bottom-right) (59). Boxplots are colored by the benchmarked method. The red line indicates the median null MSE when generating cell-type proportions randomly. If a red line is not provided, then the median null MSE is above the scale provided on the *Y*-axis.

We also investigated how the number of component compartments affects the performance of all six reference-free methods. We generated another set of *in-silico* mixed targeted panels (500 genes) using 2 (mammary tissue and lymphocytes), 3 (mammary, lymphocytes, fibroblasts), and 4 (mammary, fibroblasts, lymphocytes, and adipose) and applied all six methods to estimate the compartment proportions. **Figure 2B** provides boxplots of the MSE across 25 simulated targeted panels using *DeCompress* and the other 5 benchmarked methods. For all 6 methods, the median MSE for these datasets remained similar as the number of compartments increased, though the range in the MSE decreases considerably. In particular, the performance of *DeconICA* increases considerably as more compartments were used for mixing, as mentioned in its documentation (24). Here again, we found that *DeCompress* gave the smallest median MSE between the true and estimated cell proportions. In total, results from these *in-silico* mixing experiments show both the accuracy and precision of *DeCompress* in estimated compartment proportions.

The four cell types we used for the above analyses simulated bulk mammary tissue but contained compartments with highly correlated gene expression profiles (**Supplemental Figure 2A**). We recreated the *in-silico* mixing experiments with four compartments with minimal correlations: mammary tissue, pancreas, pituitary gland, and whole blood (**Supplemental Figure 2A**). In mixtures with these tissues, we found that *DeCompress* also outperformed the reference-free methods, with a clear decrease in median MSE as the number of genes on the simulated targeted panels are increased (**Supplemental Figure 2B**). This trend between MSE and number of genes in this setting provides some evidence that dissimilar compartments may be easier to deconvolve with more genes on the targeted panel.

#### Publicly available datasets

Although *in-silico* mixing experiments with GTEx data showed strong performance of *DeCompress*, we sought to benchmark *DeCompress* against reference-free methods in previously published datasets with known compartment mixture proportions. We downloaded expression data from a breast cancer cell-line mixture (RNA-seq) (23), rat brain, lung, and liver cell-line mixture (microarray) (11), prostate tumor with compartment proportions estimated with laser-capture microdissection (microarray) (58), and lung adenocarcinoma cell-line mixture (RNA-seq) (59) and generated pseudo-targeted panels with 200, 500, and 800 genes (see **Methods**). For the rat mixture dataset, we trained the compression sensing model on a randomly selected training split with added noise to simulate a batch effect between the training and targeted panel; for the other three cancer-related datasets, reference RNA-seq data was downloaded from The Cancer Genome Atlas (TCGA) (2). We then performed semi-reference-free deconvolution in these datasets using *DeCompress* and the reference-free methods.

Overall, *DeCompress* showed the lowest MSE across all three datasets, in comparison to the other reference-free methods (**Figure 2C**). The patterns observed in the GTEx results are evident in these real datasets, as well. As the number of genes in the targeted panel increases, the range in the distribution of MSEs decreases. Deconvolution using *Linseed* gave variable performance across datasets (high variability in model performance), with very small ranges in MSEs in the rat microarray and lung adenocarcinoma datasets while highly variable MSEs in the breast cancer and prostate cancer datasets. We do not present *DeconICA* in these comparisons due to its large errors across all datasets (see **Supplemental Figure S3** for comparisons to *DeconICA*). Specific to *DeCompress*, we assessed the performance of different deconvolution methods (4 reference-free methods and *unmix* from the *DESeq2* package (57)) on the DeCompressed expression matrix for the breast, prostate, and lung cancer datasets (**Supplemental Figure S4**). We found that *unmix* gives accurate estimates of compartment proportions in the breast cancer and prostate tumor datasets, where the component compartments are like those in bulk tumors. However, in the case of the lung adenocarcinoma mixing dataset (mixture of two lung cancer cell lines), *unmix* does not consistently outperform the reference-free methods, perhaps owing to a dissimilarity between the lung adenocarcinoma mixture dataset and TCGA-LUAD reference dataset. We lastly investigated a scenario where the reference and target assays measure different bulk tissue. Using the breast cancer cell-line mixtures pseudo-targets and a TCGA-LUAD reference, *DeCompress* estimated compartment proportions with larger errors, such that the distribution of MSEs intersect with a null distribution of MSEs from randomly generated compartment proportion matrices (**Supplemental Figure S5**).

#### Carolina Breast Cancer Study (CBCS) expression

We finally benchmarked *DeCompress* against the other 5 reference-free deconvolution methods in breast tumor expression data from the Carolina Breast Cancer Study (CBCS) (43, 55) on 406 breast cancer-related genes on 1,199 samples. We used RNA-seq breast tumor expression from TCGA to train the compression matrix for deconvolution in CBCS using *DeCompress*; 393 of the 406 genes on the CBCS panel were measured in TCGA-BRCA. For validation, a study pathologist trained a computational algorithm to estimate compartment proportions using 148 tumor microarrays (TMAs) (89). We treat these estimated compartment proportions for epithelial tumor, adipose, stroma, and immune infiltrate as a “gold standard.”

To determine whether the DeCompressed expression matrix accurately predicts expression for samples in the target, we split the 393 genes into 5 groups and trained TCGA-based predictive models of genes in each group using those in the other four. Overall, in-sample cross-validation prediction per-sample in TCGA is strong (median adjusted *R*^2^ = 0.53), with a drop-off in out-sample performance in CBCS (median adjusted *R*^2^ = 0.38), shown in **Figure 3A**. We also trained models stratified by estrogen-receptor (ER) status, a major, biologically-relevant classification in breast tumors (90, 91). These ER-specific models showed slightly better out-sample performance (median adjusted *R*^2^ = 0.34), though in-sample performance was similar to overall models with the same median *R*^2^ (**Figure 3B**). Next, as in the GTEx mixing simulations and the 4 published datasets, *DeCompress* recapitulated true compartment proportions with the minimum error (**Figure 3B**), approximately 33% less error than *TOAST + NMF*, the second-most accurate method. To provide some context to the magnitude of these errors, we randomly generated 10,000 compartment proportion matrices for 148 samples and 4 compartments. The mean MSE is provided in **Figure 3B**, showing that 2 of the 5 benchmarked methods (*CellDistinguisher* and *DeconICA*) exceeded this randomly generated null MSE value. We also observed that correlations between true and *DeCompress*-estimated compartment proportions are positive and significantly non-zero for three of four compartment components (**Figure 3C**). Unlike those from *TOAST + NMF*, *DeCompress* estimates of compartment-specific compartment proportions were positively correlated with the truth (**Supplemental Figure S6**).

**Figure 3:**
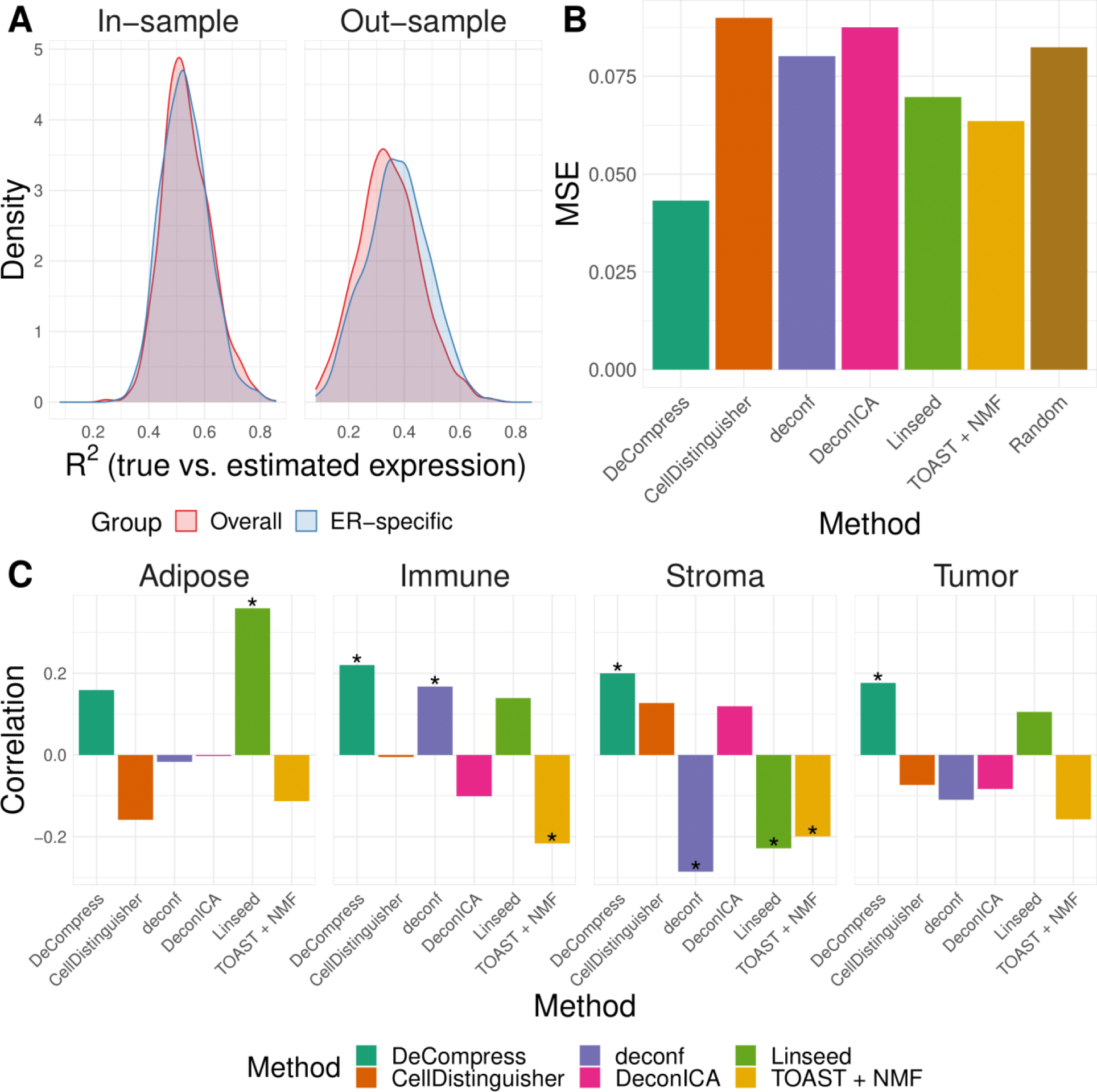
Benchmarking results with Carolina Breast Cancer Study expression data. **(A)** Kernel density plots of predicted adjusted *R*^2^ per-sample in in-sample TCGA prediction (left) through cross-validation and out-sample prediction in CBCS (right), colored by overall and ER-specific models. **(B)** MSE (*Y*-axis) between true and estimated cell-type proportions in CBCS across all methods (*X*-axis). Random indicates the mean MSE over 10,000 randomly generated cell-type proportion matrices. **(C)** Spearman correlations (*Y*-axis) between compartment-wise true and estimated proportions across all benchmarked methods (*X*-axis). Correlations marked with a star are significantly different from 0 at *P* < 0.05.

#### Comparison of computational speed

The computational cost of *DeCompress* is high, owing primarily to training the compressed sensing models. Non-linear estimation of the columns of the compression matrix is particularly slow (**Supplemental Figure S7**). In practice, we recommend running an elastic net method (LASSO, elastic net, or ridge regression) which are both faster (**Supplemental Figure S7**) and give larger cross-validation *R*^2^ (**Supplemental Figure S1**). The median cross-validation *R*^2^ for elastic net and ridge regression is approximately 16% larger than least angle regression and LASSO, and nearly 25% larger than the non-linear optimization methods. Using CBCS data with 1,199 samples and 406 genes, we ran all benchmarked deconvolution methods 25 times and recorded the total runtimes (**Supplemental Figure S8**). For *DeCompress*, we used TCGA-BRCA data with 1,212 samples as the reference. As shown in **Supplemental Figure S8**, running *DeCompress* in serial (approximately 62 minutes) takes around 40 times longer than the slowest reference-free deconvolution method (*TOAST + NMF*, approximately 1.5 minutes), though *DeCompress* is comparable in runtime to *TOAST + NMF* if run in parallel with enough workers (approximately 2.6 minutes). These computations were conducted on a high-performance cluster (RedHat Linux operating system) with 25 GB of RAM.

### Applications of DeCompress in the Carolina Breast Cancer Study

Given the strong performance of DeCompress in benchmarking experiments, we estimated compartment proportions for 1,199 subjects in CBCS with transcriptomic data assayed with NanoString nCounter. Using TCGA breast cancer (TCGA-BRCA) expression as a training set, we iteratively searched for cell type-specific features (25) (Step 1 in **Figure 1**) and included canonical compartment markers for guidance using *a priori* knowledge (30, 92, 93) (see **Methods**). After expanding the targeted CBCS expression to these genes, we estimated proportions for 5 compartments. As reference-free methods output proportions for agnostic compartments, identifying approximate descriptors for compartments is often difficult. Here, we first outline a framework for assigning modular identifiers for compartments identified by *DeCompress*, guided by compartment-specific gene signatures. Then, we assess performance of using compartment-specific proportions in downstream analyses of breast cancer outcomes and gene regulation.

#### Identifying approximate modules for DeCompress-estimated compartments

We leveraged compartment-specific gene signatures to annotate each compartment with modular identifiers. First, we computed Spearman correlations between the compartment-specific gene expression profiles and median tissue-specific expression profiles from GTEx (53, 54) and single cell RNA-seq profiles of MCF7 breast cancer cells (94) (**Figure 4A**). Here, we find that Compartment 4 (C4) shows strong positive correlations with fibroblasts, lymphocytes, multiple collagenous organs (such as blood vessels, skin, bladder, vagina, and uterus (95–97)), and MCF7 cells. We hypothesize that strong correlation with lymphocytes reflects tumor-infiltrating lymphocytes. The C3 gene signature was significantly correlated with expression profiles of secretory organs (salivary glands, pancreas, liver) and contained a strong marker of HER2-enriched breast cancer (*ERBB2*) (98).

**Figure 4:**
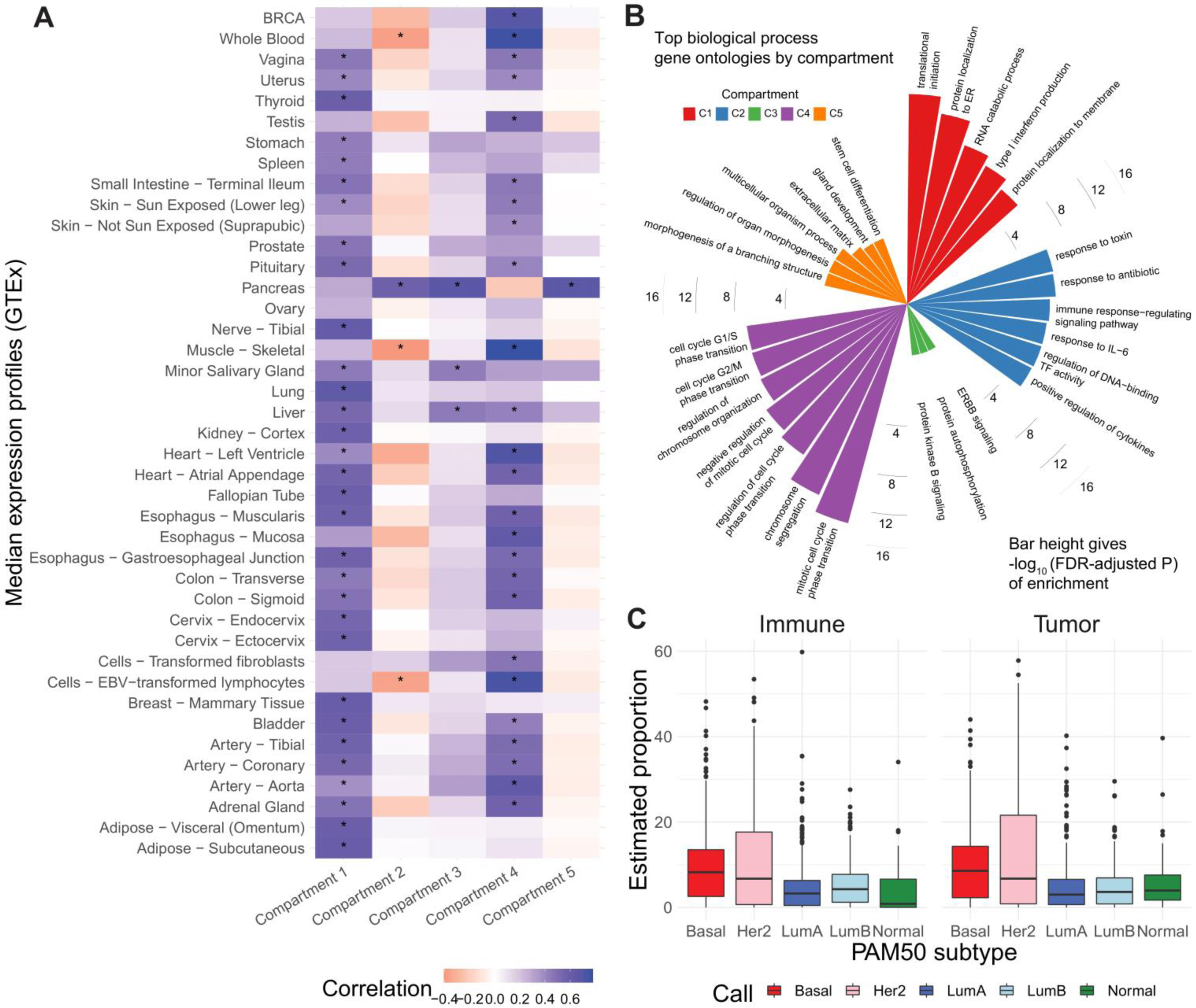
Identification of Decompress-estimated compartments. **(A)** Heatmap of Pearson correlations between compartment-specific gene signatures (*X*-axis) and GTEx median expression profiles and MCF7 single-cell profiles (*Y*-axis). Significant correlations at nominal *P* < 0.01 are indicated with an asterisk. **(B)** Bar plot of −log_10_ *FDR*-adjusted *P*-values for top gene ontologies (*Y*-axis) enriched in compartment-specific gene signatures. **(C)** Boxplots of estimated immune (left) and tumor (C3 + C4 compartments, right) proportions (*Y*-axis) across PAM50 molecular subtypes (*X*-axis)

We conducted over-representation analysis (ORA) (65) of gene signatures for all five compartments, revealing cell cycle regulation ontologies for C4 that are consistent with the hypothesis generated from GTEx profiles at FDR-adjusted *P < 0.05* (**Figure 4B**). We conducted gene set enrichment analysis (GSEA) for the C4 gene signature (99), revealing significant enrichments for cell differentiation and development process ontologies (**Supplemental Figure S9**). ORA analysis also assigned immune-related ontologies to the C2 gene signatures at FDR-adjusted *P* < 0.05 and ERBB signaling to C4, though this enrichment did not achieve statistical significance. C1 and C5 gene signatures were not enriched for ontologies that allowed for conclusive compartment assignment, showing catabolic, morphogenic, and extracellular process ontologies (**Figure 4B**). From these results, we hypothesized that C3 and C4 resembled epithelial tumor cells, C2 an immune compartment (possibly excluding lymphocytes that may infiltrate tumors), and C1 and C5 presumptively stromal and/or mammary tissue.

Distributions of the hypothesized immune (C2) and tumor (C3 + C4 proportions) revealed significant differences across PAM50 molecular subtypes (**Figure 4C**; Kruskal-Wallis test of differences with *P* < 2.2 × 10^−16^) (69). These trends across subtypes were consistent with evidence that Basal-like and HER2-enriched subtypes had the largest proportions of estimated tumor and immune compartments, while Luminal A, Luminal B, and Normal-like subtypes showed lower proportions (43, 69, 100). Furthermore, we found strong differences in C4 and total tumor compartment estimates across race (**Supplemental Figure S10A**). C3 and C4 also have strong correlations with ER- (estrogen receptor) and HER2-scores, gene-expression based continuous variables that indicate clinical subtypes based on *ESR1* and *ERBB2* gene modules (**Supplemental Figure S10B**); however, none of the C3, C4, immune, or tumor compartment estimates showed significant differences across clinical ER status determined by immunohistochemistry (**Supplemental Figure S10C**). We considered the incorporation of estimates of compartment proportions in building models of breast cancer survival (**Supplemental Results** and **Supplemental Table S3**).

#### Incorporating compartment proportions into eQTL models detects more tissue-specific gene regulators

We investigated how incorporating estimated compartment proportions affect *cis*-expression quantitative trait loci (*cis*-eQTL) mapping in breast tumors, a common application of deconvolution methods in assessing sources of variation in gene regulation (9, 101). In previous eQTL studies using CBCS expression, several bulk breast tumor *cis*-eGenes (i.e. the gene of interest in an eQTL association between SNP and gene expression) were found in healthy mammary, subcutaneous adipose, or lymphocytes from GTEx (10). We included *DeCompress* proportion estimates for the tumor (C3 + C4 estimates) and immune (C2) compartments in a race-stratified, genetic ancestry-adjusted *cis*-eQTL interaction model (see **Methods**), as proposed by Geeleher *et al* and Westra *et al* (8, 9). We found that sets of compartment-specific *cis*-eGenes generally had few intersections with bulk *cis*-eGenes (**Figure 5A**), though we detected more *cis*-eQTLs with the immune- and tumor-specific interaction models (**Supplemental Figure S11**). At FDR-adjusted *P* < 0.05, of 209 immune-specific *cis*-eGenes identified in women of European ancestry (EA), 7 were also mapped in the bulk models (with no compartment proportion covariates), and no tumor-specific *cis*-eGenes were identified with the bulk models. Similarly, at FDR-adjusted *P* < 0.05, in women of African ancestry (AA), 27 of 331 and 9 of 124 *cis*-eGenes identified with the immune- and tumor-compartment interaction models were also mapped with the bulk models, respectively. Manhattan plots for *cis*-eQTLs across the whole genome across bulk, tumor, and immune show the differences in eQTL architecture in these compartment-specific eQTL mappings in EA and AA samples (**Supplemental Figures S12** and **S13**, respectively). Furthermore, we generally detected more *cis*-eQTLs at FDR-adjusted *P* < 0.05 with the immune-specific interactions than the bulk and tumor-specific interactions (EA: 565 bulk *cis*-eQTLs, 65 tumor *cis*-eQTLs, 8927 immune *cis*-eQTLs; AA: 237 bulk *cis*-eQTLs, 449 tumor *cis*-eQTLs, 7676 immune *cis*-eQTLs; **Supplemental Figure S11**). All eQTLs with FDR-adjusted *P* < 0.05 are provided in **Supplemental Data** (https://github.com/bhattacharya-a-bt/DeCompress_supplement) (102).

**Figure 5:**
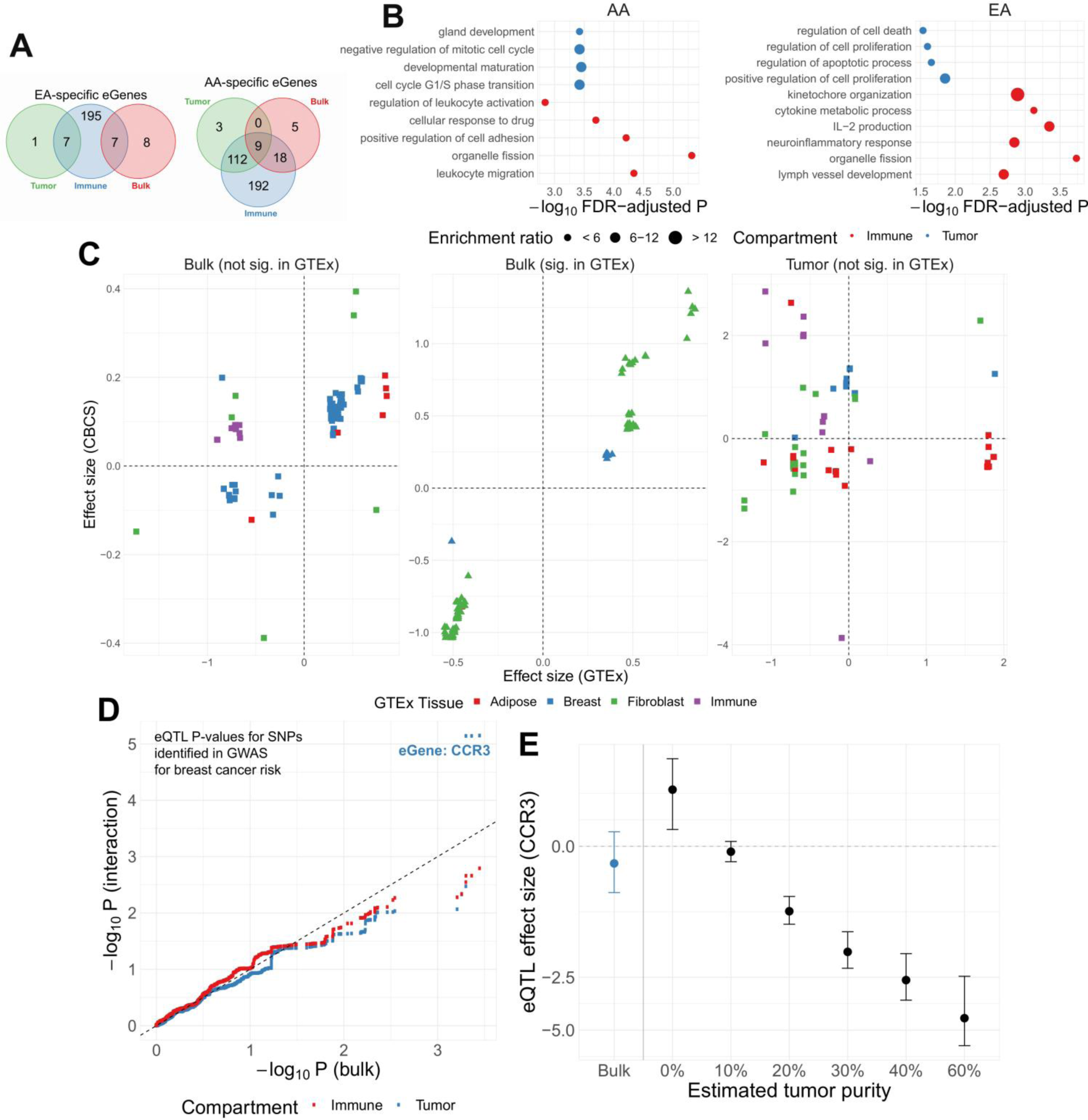
Compartment-specific cis-eQTL mapping in the Carolina Breast Cancer Study. **(A)** Venn diagram of bulk, tumor-, and immune-specific *cis*-eGenes identified European-ancestry (left) and African-ancestry samples (right) in CBCS. **(B)** Enrichment analysis of immune- (red) and tumor-specific (blue) cis-eGenes in CBCS plotting the *−log*_10_ *P*-value of enrichment (*X*-axis) and description of gene ontologies (*Y*-axis). The size of the point represents the relative enrichment ratio for the given ontology. **(C)** Scatterplots of GTEx (*X*-axis) and CBCS effect size (*Y*-axis) for significant CBCS *cis*-eQTLs that were mapped in GTEx. Each point is colored by the GTEx tissue in which the cis-eQTL has the lowest *P*-value. Reference dotted lines for the *X*- and *Y*-axes are provided. **(D)** For risk variants from GWAS for breast cancer from iCOGs (86–88), scatterplot of *−log*_10_ *P*-values of bulk (*X*-axis) and compartment-specific *cis*-eQTLs (*Y*-axis), colored blue for tumor- and red for immune-specific models. A 45-degree reference line is provided. In the top right corner, 3 tumor-specific *cis*-eQTLs are labeled with the eGene *CCR3* as they are significant at FDR-adjusted *P* < 0.05. **(E)** Tumor-specific eQTL effect sizes and 95% confidence intervals (*Y*-axis) for rs56387622 on *CCR3* expression across various estimates of tumor purity. The eQTL effect size from the bulk model is given in blue.

We analyzed the sets of EA and AA tumor- and immune-specific eGenes in CBCS with ORA analysis for biological processes (**Figure 5B**). We found that, in general, these sets of eGenes were concordant with the compartment in which they were mapped. All at FDR-adjusted *P* < 0.05, AA tumor-specific eGenes showed enrichment for cell cycle and developmental ontologies, while immune-specific eGenes were enriched for leukocyte activation and migration and response to drug pathways. Similarly, EA tumor-specific eGenes showed enrichments for cell death and proliferation ontologies, and immune-specific eGenes showed cytokine and lymph vessel-associated processes. We then cross-referenced bulk and tumor-specific *cis*-eGenes found in the CBCS EA sample with *cis*-eGenes detected in healthy tissues from GTEx: mammary tissue, fibroblasts, lymphocytes, and adipose (see **Methods**), similar to previous pan-cancer germline eQTL analyses (10, 103). We attributed several of the bulk *cis*-eGenes to healthy GTEx tissue (all but 2), but tumor-specific *cis*-eGenes were less enriched in healthy tissues (**Supplemental Figure S14**). We compared the *cis*-eQTL effect sizes for significant CBCS *cis*-eSNPs found in GTEx. As shown in **Figure 5C**, 98 of 220 bulk *cis*-eQTLs detected in CBCS that were also found in GTEx were mapped in healthy tissue, with strong positive correlation between effect sizes (Spearman *ρ* = 0.93). The remaining 122 eQTLs that could not be detected in healthy GTEx tissue contained some discordance in the direction of effects, though correlations between these effect sizes were also high (*ρ* = 0.71). In contrast, we were unable to detect any of the CBCS tumor-specific *cis*-eQTLs in as significant eQTLs in GTEx healthy tissue, and the correlation of these effect sizes across CBCS and GTEx was poor (Spearman *ρ* = −0.07). These results suggest that this compartment-specific eQTL mapping, especially those that are tumor-specific, identified eQTLs that are not enriched for eQTLs from healthy tissue.

To evaluate any overlap of compartment-specific eQTLs with SNPs implicated with breast cancer risk, we extracted 932 risk-associated SNPs in women of European ancestry from iCOGS (86–88) at FDR-adjusted *P* < 0.05 that were available on the CBCS OncoArray panel (71). **Figure 5D** shows the raw − log_10_ *P*-values of the association of these SNPs with their top cis-eGenes in the bulk and tumor- and immune-specific interaction models. In large part, none of these eQTLs reached FDR-adjusted *P* < 0.05, except for three *cis*-eQTLs, with their strengths of association favoring the bulk eQTLs. However, we detected 3 tumor-specific EA *cis*-eQTLs in near-perfect linkage disequilibrium of *r*^2^ ≥ 0.99 (strongest association with rs56387622) with chemokine receptor *CCR3*, a gene whose expression was previously found to be associated with breast cancer outcomes in luminal-like subtypes (104, 105). As estimated tumor purity increases, the cancer risk allele C at rs56387622 has a consistently strong negative effect on *CCR3* expression (**Figure 5E**). We find that *CCR3* expression is insignificantly different across tumor stage and ER status but is significantly different across PAM50 molecular subtype (**Supplemental Figure S15**). In sum, results from our *cis*-eQTL analysis show the advantage of including *DeCompress*-estimated compartment proportions in downstream genomic analyses to identify compartment-specific associations that may be relevant in disease pathways.

## DISCUSSION

Here, we presented *DeCompress*, a semi-reference-free deconvolution method catered towards targeted expression panels that are commonly used for archived tissue in clinical and academic settings (3, 35). Unlike traditional reference-based methods that require compartment-specific expression profiles, *DeCompress* requires only a reference RNA-seq or microarray dataset on similar bulk tissue to train a compressed sensing model that projects the targeted panel into a larger feature space for deconvolution. Such reference datasets are much more widely available than compartment-specific expression on the same targeted panel. We benchmarked *DeCompress* against reference-free methods (20, 22, 24–26) using *in-silico* GTEx mixing experiments (53, 54), 4 published datasets with known compartment proportions (11, 23, 58, 59), and a large, heterogeneous NanoString nCounter dataset from the CBCS (43, 55). In these analyses, we showed that *DeCompress* recapitulated true compartment proportions with the minimum error and the strongest compartment-specific positive correlations, especially when the reference dataset is properly aligned with the tissue assayed in the target. We tested the performance of *DeCompress* by incorporating compartment estimates in eQTL mapping to reveal immune- and tumor-compartment-specific breast cancer eQTLs.

While *DeCompress* has several important strengths, it has some limitations. First, *DeCompress* has a high computational cost, owing mainly to its lengthy compressed sensing training step. We recommend running mainly linear optimization methods in this step and have implemented parallelization options to bring computation time on par with the iterative framework proposed in TOAST (25). However, *DeCompress* estimates compartment proportions both accurately and precisely, compared to other reference-free methods, and provides a strong computational alternative that is much faster than costly lab-based measurement of composition. Second, *DeCompress*, as a semi-reference-free method, shares the limitations of reference-based methods – namely concerns with the proper selection of a reference dataset. As seen in the lung adenocarcinoma example, where TCGA-LUAD data was not an accurate reflection of a mixture of adenocarcinoma cell-lines, *DeCompress* performance has slightly lower performance than datasets properly matched to their references. Yet, in this setting, *DeCompress* performance was on par with that of the other reference-free methods that do not use a misaligned reference. Lastly, also in common with reference-free methods, the compression model may also be sensitive to phenotypic variation in the reference, as evidenced by the increase in out-sample prediction *R*^2^ in ER-specific models compared to overall models in CBCS. This specificity may be leveraged to train more accurate models by using more than one reference dataset to reflect clinical or biological heterogeneity in the targeted panel. Researchers may employ more systematic methods of assessing the similarity of the reference and target datasets, like measuring the distance between the two matrices (i.e. norms based on the singular values of matrices) or comparing the correlation structure of overlapping genes in the feature spaces of the reference and target. These evaluations will help with selecting a proper reference for a targeted panel to be deconvolved using *DeCompress*.

*DeCompress* also shares some challenges with reference-free deconvolution methods, such as the selection of an appropriate number of compartments. Previous groups have emphasized reliance on *a priori* knowledge for deconvolving well-studied tissues, such as blood and brain (106, 107). However, diseased tissues, like bulk cancerous tumors, especially in understudied subtypes or populations, are more difficult to deconvolve due to the similarity between compartments, many of which may be rare or reflect transient cell states (30, 91, 108, 109). For this reason, we included several data-driven approaches in estimating the number of compartments from variation in the gene expression and recommended applying prior domain knowledge about the tissue of interest. It is also important to carefully consider the gene module-based annotations for the unidentified estimated compartments, especially in bulk tissue where traditional ideas of compartments are inapplicable (29). Several previous reference-free methods have leveraged *in vitro* mixtures of highly distinct cell lines in training and testing previous reference-free deconvolution methods (11, 22), namely the rat cell line mixture (GSE19830) (11). Though this dataset is easy to deconvolve and thus useful in testing methodology, the extreme differences in gene expression between these three tissue types renders this dataset sub-optimal for methods benchmarking. Furthermore, assigning estimated compartments to known tissues in this dataset is straightforward and does not capture the difficulty of this task in typical deconvolution applications. Instead, our applications in breast cancer expression with CBCS provided such a difficult statistical challenge. Our outlined approach of first comparing compartment-specific gene signatures to known tissue profiles from GTEx or single-cell profiles, then analyzing these signatures with ORA or GSEA, and lastly checking hypotheses against known biological trends provides a structured framework for addressing the compartment identification problem.

Our downstream eQTL analysis in CBCS breast tumor expression also provided some insight into gene regulation, similar to recent work into deconvolving immune subpopulation eQTL signals from bulk blood eQTLs (101). In breast cancer, Geeleher *et al* previously showed that a similarly implemented interaction eQTL model gave better mapping of compartment-specific eQTLs (8, 9). Our results are consistent with this finding, especially since tumor- and immune-specific eGenes were enriched for commonly associated ontologies. However, unlike Geeleher et al, we generally detected a larger number of immune- and tumor-specific eQTLs and eGenes than in the bulk, unadjusted models. We believe that this larger number of compartment-specific eGenes may be due to the specificity of the genes assayed by the CBCS targeted panel. As the panel included 406 genes, all previously implicated in breast cancer pathogenesis, proliferation, or response (10, 43, 110), the interaction model will detect SNPs that have large effects on compartment-specific genes. The interaction term is interpreted as the difference in eQTL effect sizes between samples of 0% and 100% of the given compartment; accordingly, for genes implicated in specific breast cancer pathways, we expect to see large differences in compartment-specific eQTL effects (111–113). Though this interaction model is straight-forward in its interpretation for the tumor compartment (i.e. a sample of 100% tumor cells versus 100% tumor-associated normal cells), this interpretation may be tenuous for less well-defined compartments, like an immune compartment that includes several different immune cells. This interaction term’s effect size may also be inflated for compartment estimates that have low mean and high variance across the samples. In addition, we did not consider *trans*-acting eQTLs that are often attributed to compartment heterogeneity, though we believe that methods employing mediation or cross-condition analysis can be integrated with compartment estimates to map compartment-specific *trans*-eQTLs relevant in breast cancer (114–116).

Relevant to risk and proliferation of breast cancer, we detected a locus of *cis*-eSNPs associated with expression of *CCR3* (C-C chemokine receptor type 3) that were GWAS-identified risk SNPs (86–88) but were not significantly associated with *CCR3* expression using the bulk models and were not detected in GTEx. If one or more causal SNPs in this genomic region affects *CCR3* expression only in cancer cells and the effect on *CCR3* expression is the main mechanism by which the locus predisposes individuals to breast cancer, we can hypothesize that an earlier perturbation in the development of cancer (e.g. transcription factor or microRNA activation) may cause this SNP’s tumorigenic effect. Given this perturbation in precancerous mammary cells, individuals with the risk allele would convey the tumorigenic effects of decreased *CCR3* expression. It has been previously shown that increased peritumoral *CCR3* expression is associated with improved survival times in luminal-like breast cancers (104, 105). The CCR3 receptor has been shown to be the primary binding site of CCL11 (eotaxin-1), an eosinophil-selective chemoattractant cytokine (117, 118), and accordingly CCR3 antagonism prohibited chemotaxis of basophils and eosinophils, a phenomenon observed in breast cancer activation and proliferation (119, 120). Without *DeCompress* and the incorporation of estimated compartment proportions in the eQTL model, this association between eSNP and *CCR3* expression would not have been detected in this dataset (121).

*DeCompress*, our semi-reference-free deconvolution method, provides a powerful method to estimate compartment-specific proportions for targeted expression panels that have a limited number of genes and only requires RNA-seq or microarray expression from a similar bulk tissue. Our method’s estimates recapitulate known compartments with less error than reference-free methods and provides compartments that are biologically relevant, even in complex tissues like bulk breast tumors. We provide examples of using these estimated compartment proportions in downstream studies of outcomes and eQTL analysis. Given the wide applications of reference-free deconvolution, the popularity of targeted panels in both academic and clinical settings, and increasing need for analyzing heterogeneous and dynamic tissues, we anticipate creative implementations of *DeCompress* to give further insight into expression variation in complex diseases.

## Supporting information

Additional File 1: Supplemental Methods, Results, Tables, and Figures

## DATA AVAILABILITY

The *DeCompress* package is available as R software on GitHub: https://github.com/bhattacharya-a-bt/DeCompress. Sample code for replication and results from the eQTL analysis are provided: https://github.com/bhattacharya-a-bt/DeCompress_supplement (102). CBCS expression data is publicly available at GSE148426. CBCS genotype datasets analyzed in this study are not publicly available as many CBCS patients are still being followed and accordingly is considered sensitive; the data is available from M.A.T upon reasonable request. GTEx median expression profiles are available from dbGAP accession number phs000424.v7.p2. Data from the published mixture experiments are available from GEO: GSE19830, GSE123604, GSE97284, and GSE64098. Single-cell expression profiles of MCF7 cells were obtained from GSE52716. Expression data from The Cancer Genome Atlas is available from the Broad GDAC Firehose repository (https://gdac.broadinstitute.org/) with accession number phs000178.v11.p8.

## SUPPLEMENTARY DATA

Additional File 1: Supplemental Methods, Results, Tables, and Figures

## ACKNOWLEDGEMENTS

We think the Carolina Breast Cancer Study participants and volunteers. We also thank Katie Hoadley, Linnea Olsson, Chuck Perou, Luca Pinello, and Naim Rashid for valuable discussion during the research process. We thank Erin Kirk and Jessica Tse for their invaluable support during the research process. We thank the DCEG Cancer Genomics Research Laboratory and acknowledge the support from Stephen Chanock, Rose Yang, Mereditch Yeager, Belynda Hicks, and Bin Zhu.

The Genotype-Tissue Expression (GTEx) Project was supported by the Common Fund of the Office of the Director of the National Institutes of Health, and by NCI, NHGRI, NHLBI, NIDA, NIMH, and NINDS. The data used for the analyses described in this manuscript were obtained from: the GTEx Portal (https://storage.googleapis.com/gtex_analysis_v7/rna_seq_data/GTEx_Analysis_2016-01-15_v7_RNASeQCv1.1.8_gene_median_tpm.gct.gz) with dbGaP accession number phs000424.v7.p2 on 05/14/20.

