## Additional File 1: Supplemental Methods, Results, Tables, and Figures for "DeCompress: tissue compartment deconvolution of targeted mRNA expression panels using compressed sensing"

Supplemental Materials

#### 1 Supplemental Methods

##### 1.1 DeCompress algorithm

DeCompress takes in two expression matrices from similar bulk tissue as inputs: the *target* matrix $\mathbf{T}$, an $n \times k$ matrix from a targeted panel of gene expression, and the *reference* matrix $\mathbf{R}$, an $N \times K$ matrix from an RNA-seq or microarray panel, such that $K > k$. For a user-defined $c$ cell-types, DeCompress outputs $\hat{\mathbf{S}}$, a $c \times K'$ matrix of cell-type specific expression profiles and $\hat{\mathbf{P}}$, a $c \times n$ matrix of cell-type proportions. The method follows three general steps, as detailed in **Figure 1**: (1) selection of approximate cell-type specific genes, (2) compressed sensing to expand the feature space of $\mathbf{T}$, and (3) ensemble reference-free deconvolution on expanded expression dataset. DeCompress is freely available as an R package on Github (`https://github.com/bhattacharya-a-bt/DeCompress`).

###### 1.1.1 Selection of cell-type specific genes

The first step of DeCompress is to find a set of $K' < K$ genes that are representative of the different cell types that comprise the bulk tissue. These $K'$ genes, called the cell-type specific (CTS) genes, can be supplied by the user if prior gene signatures can be applied. If any such gene signatures are not available, DeCompress borrows methods from two previous reference-free deconvolution methods to select a parsimonious gene set.

We include methods from Zaitsev *et al.*'s LINear Subspace identification for gene Expression Deconvolution (LINSEED) method[1] that assumes mutual linearity (i.e. $y_1 = ky_2$, where $y_1$ and $y_2$ are the expressions of gene 1 and gene 2, respectively) between cell-type specific genes to generate gene signatures. Briefly, LINSEED transforms the gene expression space to form a $c$-vertex simplex, where each vertex represents a distinct cluster of mutually linear genes corresponding to a cell type. The algorithm then picks the closest genes to each vertex to represent a cell-type specific gene signature[1]. We also include Li and Wu's feature selection method, TOols for the Analysis of heterogeneouS Tissues (TOAST)[2], which iteratively searches for cell type-specific genes and performs reference-free estimation at each step. TOAST uses a novel hypothesis testing framework to conduct cross-cell type differential analysis and identify gene signatures[2].

###### 1.1.2 Compressed sensing framework

After a suitable set of $K'$ CTS genes are determined, we take the $K'$ corresponding columns of $\mathbf{R}$ to form $\mathbf{R'}_{N\times K'}$ and the $k$ genes corresponding to columns in $\mathbf{T}$ to form $\mathbf{R^{(k)}_{N\times k}}$. Consider the following matrix equation, where $\mathbf{\Phi}$ is a $k \times K'$ compression matrix that projects $\mathbf{R^{(k)}}$ to $\mathbf{R}'$:

$$\mathbf{R}'_{N\times K'} = \mathbf{R^{(k)}_{N\times k}\Phi_{k\times K'}} \tag{1}$$

We can break down Equation 1 into a system of equations. For the $i$th column of $\mathbf{R}'$, denoted $r'_i$, we wish to find a $k$-length sparse vector $\phi_i,\ 1 \leq i \leq K'$ such that

$$r'_i = \mathbf{R^{(k)}_{N\times k}}\phi_i. \tag{2}$$

We estimate $\hat{\phi}_i$ with the following optimization methods: least angle regression (using R package lars)[3], elastic net with elastic net mixture penalty $\alpha \in \{0, 0.5, 1\}$ (using the R package glmnet)[4], and $l_1$,$l_2$, and total variation $l_1$ (TV-L1) non-linear optimization (using R package R1magic)[5–8]. Functions in DeCompress allow the user to select any to all of these optimization methods and picks the best method through 5-fold cross-validation.

Especially when $N$ is sufficiently large, non-linear optimization is computationally expensive (see comparison of run times in **Supplemental Figure S6**). We implement parallelization across columns of $\mathbf{R}'$ using the future package in R[9] and recommend linear optimization methods as they are faster and give generally similar prediction (Supplemental Figure S1).

###### 1.1.3 Optimization methods for compressed sensing

Compressed sensing in DeCompress aims to estimate the $k \times K'$ compression matrix $\mathbf{\Phi}$ in the equation:

$$\mathbf{R}'_{N\times K'} = \mathbf{R^{(k)}_{N\times k}\Phi_{k\times K'}}. \tag{3}$$

We convert this into a system of equations. For the $i$th column of $\mathbf{R}'$, denoted $r'_i$, we wish to find a $k$-length sparse vector $\phi_i,\ 1 \leq i \leq K'$ such that

$$r'_i = \mathbf{R^{(k)}_{N\times k}}\phi_i. \tag{4}$$

DeCompress implements several regularized regression or optimization methods to estimate $\hat{\phi}_i$:

- *Elastic net*[4] finds

$$\hat{\phi}_i = \arg\min_{\phi_i} \left\{ \|r_i' - \mathbf{R}^{(\mathbf{k})}\phi_i\|_2^2 + \lambda \left[ \frac{(1-\alpha)}{2} \|\phi_i\|_2^2 + \alpha\|\beta\|_1 \right] \right\}. \tag{5}$$

We have implemented $\alpha \in \{0, 0.5, 1\}$, where $\alpha = 0$ represents ridge regression with no sparsification of $\phi_i$ and $\alpha = 1$ represents traditional Lasso[10]. This optimization is carried out in DeCompress with the glmnet package[4].

- *Least angle regression* (LARS) minimizes the Lasso objective function in Expression 5 that speeds ups stage-wise forward selection. The algorithm starts with all elements of $\phi_i$ equal to zero and finds the predict most correlated with the response. The largest step possible is take in the direction of these predictor until some other predictor has as much correlation with the residual. LARS then proceeds in a direction equiangular between these two predictors until a third variable shares an equal correlation with the residual. The full mathematical justification and details are provided by Efron *et al.*[3]

- $l_1$ *non-linear optimization* solves the following optimization using the `nlm` function in R, as implemented in the R1magic package[5]:

$$\hat{\phi}_i = \arg\min_{\phi_i} \left\{ \sum_{i=1}^{N} |\mathbf{R}^{(\mathbf{k})}\mathbf{T}\phi_i - r_i'|^2 + \lambda|\phi_i| \right\}, \tag{6}$$

where $\mathbf{T}$ is a $K' \times K'$ matrix of sparsity bases and $\lambda$ is a tuned penalty parameter.

- $l_2$ *non-linear optimization* solves the following optimization using the `nlm` function in R, as implemented in the R1magic package[5]:

$$\hat{\phi}_i = \arg\min_{\phi_i} \left\{ \sum_{i=1}^{N} |\mathbf{R}^{(\mathbf{k})}\mathbf{T}\phi_i - r_i'|^2 + \lambda\sqrt{|\phi_i|} \right\}, \tag{7}$$

where $\mathbf{T}$ is a $K' \times K'$ matrix of sparsity bases and $\lambda$ is a tuned penalty parameter.

- *total-variation* $l_1$ *non-linear optimization* solves the following optimization using the `nlm` function in R, as implemented in the R1magic package[5]:

$$\hat{\phi}_i = \arg\min_{\phi_i} \left\{ \sum_{i=1}^{N} \|\mathbf{R}^{(\mathbf{k})}\mathbf{T}\phi_i - r_i'\|_F^2 + \lambda TV(\phi_i) \right\}, \tag{8}$$

where $\mathbf{T}$ is a $K' \times K'$ matrix of sparsity bases, $\lambda$ is a penalty parameter, and $TV(\cdot)$ is the total-variation function, such that for a generic $n$-length vector $\nu$ with $j$th element $\nu_j$

$$TV(\nu) = \sum_{i=1}^{n-1} |\nu_i - \nu_{i+1}|.$$

##### 1.1.4 Ensemble deconvolution on expanded dataset

After the estimated compression matrix $\hat{\mathbf{\Phi}}$ is obtained, we then expand the expression matrix from the targetted panel $\mathbf{T}_{n\times k}$ into a larger features space by multiplying $\mathbf{T}$ with $\hat{\mathbf{\Phi}}$:

$$\tilde{\mathbf{T}}_{n\times K'} = \mathbf{T}_{n\times k}\hat{\mathbf{\Phi}}_{k\times K'}.$$

This expanded expression matrix $\tilde{\mathbf{T}}$, called the *decompressed* expression matrix, is then used for ensemble deconvolution. DeCompress includes multiple options for deconvolution, summarized in **Supplemental Table S1**: (1) reference-free methods, such as deconf[11], CellDistinguisher[12], TOAST with non-negative matrix factorization[2], Linseed[1], and DeconICA[13], and (2) reference-based methods using cell-type specific expression profiles from factorization of $\mathbf{R}'_{N\times K'}$, unmix from the DESeq2 package[14]. The optimal estimated cell-type proportion matrix $\hat{\mathbf{P}}$ and cell-type specific expression profiles matrix $\hat{\mathbf{S}}$ are selected from the method that best recreates $\tilde{\mathbf{T}}$ (i.e. minimizes $\|\tilde{\mathbf{T}} - \hat{\mathbf{S}}^T\hat{\mathbf{P}}\|$).

#### 1.2 In-silico GTEx mixing experiments

We downloaded median tissue-specific expression profiles from the Genotype-Tissue Expression (GTEx) Project[15,16] for mammary tissue, lymphocytes, fibroblasts, and adipose tissue. Call these median expression profiles $\mathbf{E}_{\text{profile}}$. We randomly generated a matrix of mixing proportions $\mathbf{P}$ for $n$ samples and $c \in \{2, 3, 4\}$ of the tissue types. We then generated mixed expression profiles with the following model:

$$\mathbf{E}_{\text{mixed}} = \mathbf{E}_{\text{profile}}\mathbf{P}^T.$$

We then multiplied each element of $\mathbf{E}_{\text{mixed}}$ with a randomly generated error term drawn from a Normal distribution with 0 mean and standard deviation of either 4 or 8 (low and high noise). This simulates natural perturbation to mixed expression profiles. We then randomly generated 25 simulated pseudo-targted panels each of $K \in \{200, 500, 800, 1000\}$ genes that have means and variances above the median mean and variance of all genes in the simulated genes. These simulated datasets have sample size 200. For benchmarking, in each of these simulated datasets, we selected 100 of the 200 samples as a test set for deconvolution. The other 100 samples are considered only in DeCompress deconvolution and simulated expression for all genes are kept as the reference. We added more multiplicative noise to the reference drawn from a Normal distribution with zero mean and standard deviation of 10 to simulate batch differences between the reference and target.

#### 1.3 Benchmarking in published datasets

We downloaded four datasets, as mentioned in **Methods** and summarized in **Supplemental Table S2**. Here, we detail the process of generating pseudo-targeted panels from these RNA-seq or microarray datasets. Assume the downloaded datasets are coded in the matrix $\mathbf{E}$ with $K$ rows corresponding to genes and $n$ columns corresponding to samples. We take the $K'$ genes such that the means and variances of each of these $K'$ genes are in the top 50% of means and variances of all $K$ genes. This restriction is placed on the $K'$ genes so as to not include lowly expressed genes with no variation across cell-types or other conditions. We then generated 25 pseudo-targeted panels with randomly selected 200, 500, 800, and 100 of the $K'$ genes.

### 2 Supplemental Results

#### 2.1 Incorporating estimated compartment improves outcome prediction

Next, we considered the impact of including the tumor (C3, C4, and combining C3/C4) and immune (C2) compartments in survival models. We constructed Cox models for breast-cancer specific mortality[17] with the following covariates: race, age, PAM50 molecular subtype, compartment proportion, and an interaction between subtype and compartment proportion. **Supplemental Table S3** shows hazard ratio estimates and 90% FDR-adjusted confidence intervals[18] from Cox models with the C3, C4, tumor, and immune compartments, along with comparisons to a reduced baseline model that excludes the compartment estimates and interaction terms. General relationships stay similar across the baseline and interaction models (e.g. protective hazard ratios of Luminal A subtypes in comparison to the reference Basal subtypes). We also estimated, in the C4-compartment interaction model, that increased C4 proportion was associated with shorter survival (hazard ratio 1.69, FDR-adjusted $P = 0.026$). We also compared these compartment-specific interaction models with the nested baseline model that did not contain the compartment proportions using a partial likelihood ratio test. We found that only the interaction model with the C4 proportions gave a significantly better model fit ($\chi^2 = 11.52$ on 4 degrees of freedom, $P = 0.02$). Estimated survival Kaplan-Meier curves stratified by molecular subtype and median-stratified C3 and C4 proportions showed significant differences between low and high proportion groups within molecular subtypes (**Supplemental Table S3**). Namely, we observed that the C3 high and low proportion groups only split the HER2-enriched molecular subtype based on survival outcomes, reinforcing the ERBB signaling annotations assigned to C3 in ORA analysis. However, the HER2-enriched subtype was enriched for C3-high samples (127 out of 147 samples in the C3-high group). We also found that the C4 groups split the Basal and Luminal B subtype groups, though the Basal subtype was disproportionately enriched for C4-high subjects (315 out of 339 subjects). In sum, these results illustrate that incorporating computationally-derived estimates of compartments may aid in outcome prediction.

### 3 Supplemental Tables

| Method | Summary | Implementation |
|---|---|---|
| deconf[11] | Non-negative least squares on normalized expression matrix in $\log_2$-space, seeded by initial non-negative matrix factorization. | R package CellMix[19] |
| TOAST[2] | Feature selection used in combination with iterative reference-free deconvolution. Feature selection is done using a method for cross-cell type differential analysis for data from a mixed sample[20]. | R package TOAST[2] |
| CellDistinguisher[12] | Topic modeling based on a set of input cell-type distinguishing genes. CellDistinguisher includes a method to infer distinguishing genes using the gene-gene conditional expression vectors in a space where the number of vectors and number of dimensions are both equal to the number of genes. This step relies on a large input number of genes to properly function. | R package CellDistinguisher[12] |
| Linseed[1] | Solving a convex hull problem by projecting the gene expression data and find corners using an assumption that cell-type specific genes are mutually linear. The cell-type specific expression genes are then inputted into the Digital Sorting Algorithm, a gene-signature based deconvolution method[21]. | R package linseed |
| DeconICA[13] | Deconvolution using Independent Component Analysis (ICA), a matrix factorization method for dimension reduction by projecting the expression into a space such that distributions of the data point projections on the new axes are as mutually independent as possible. | R package DeconICA[13] |
| unmix[22,14] | Non-negative least squares on the non-$\log_2$ scale with loss calculated in a variance stabilized space. This is a reference-based method, and is seeded in DeCompress using the estimated cell-type specific expression profiles estimated from the reference. | R package DESeq2[14] |

Table S1: *Summary of deconvolution methods benchmarked against or employed in DeCompress.*

| Dataset | Accession Number | Description |
| --- | --- | --- |
| In-silico GTEx mixing[15,16] | dbGAP: phs000424.v7.p2 | Median tissue-specific expression profiles were mixed at randomly generated mixing proportions to simulate targeted panels. |
| Rat tissue cell-line mixture[23] | GEO: GSE19830 | Rat brain, liver, and lung biospecimens from one animal were mixed at the cRNA homogenate level in different proportions. Expression was measured using microarray. |
| Human breast cancer cell-line mixture[24] | GEO: GSE123604 | Total mRNA was prepared from Namalwa (Burkitt's lymphoma), Hs343T (fibroblasts from mammary gland adenocarcinoma), hTERT-HME1 (normal mammary epithelial cells), and MCF7 (estrogen receptor positive breast cancer cells). Cell lines were mixed in different proportions and expression was measured using RNA-seq. |
| Human prostate tumor laser capture microdissection[25] | GEO: GSE97284 | Gene expression profiling of laser capture microdissected epithelial and stromal specimens from prostate tumors using microarray. |
| Human lung cancer cell-line mixture[26] | GEO: GSE64098 | Two lung adenocarcinoma cell lines (NCI-H1975 and HCC827) were mixed at different proportions and expression was measure using RNA-Seq. |
| Bulk breast tumors from the Carolina Breast Cancer Study[27,28] | Request GEO download token from authors | Expression from bulk breast tumors were measured using NanoString nCounter. A pathologist estimated cell-type proportions for 148 samples from tumor microarrays. |

Table S2: *Summary of datasets used in benchmarking*

| Covariate | Hazard Ratio (90% adjusted CI) | FDR-adjusted P |
|---|---|---|
| **Baseline** | | |
| *PAM50: HER2* | 1.37 (0.97,1.95) | 0.310 |
| *PAM50: LumA* | 0.55 (0.39, 0.79) | 0.041 |
| *PAM50: LumB* | 1.22 (0.86, 1.72) | 0.220 |
| *Race: White* | 0.76 (0.59, 1.00) | 0.110 |
| *Age (in 10 yrs)* | 0.84 (0.75, 0.95) | 0.072 |
| *Compartment (in 10%)* | | |
| *HER2/Compartment* | | |
| *LumA/Compartment* | | |
| *LumB/Compartment* | | |
| **C3** | | |
| *PAM50: HER2* | 1.65 (0.87, 3.11) | 0.260 |
| *PAM50: LumA* | 0.52 (0.30, 0.89) | 0.064 |
| *PAM50: LumB* | 1.15 (0.64, 2.05) | 0.782 |
| *Race: White* | 0.76 (0.57, 1.03) | 0.214 |
| *Age (in 10 yrs)* | 0.84 (0.74, 0.96) | 0.064 |
| *Compartment (in 10%)* | 1.07 (0.68, 1.68) | 0.782 |
| *HER2/Compartment* | 0.86 (0.50, 2.41) | 0.782 |
| *LumA/Compartment* | 1.12 (0.59, 2.11) | 0.782 |
| *LumB/Compartment* | 1.11 (0.51, 2.41) | 0.782 |
| **C4** | | |
| *PAM50: HER2* | 2.57 (1.42, 4.67) | 0.026 |
| *PAM50: LumA* | 0.90 (0.51, 1.60) | 0.761 |
| *PAM50: LumB* | 2.30 (1.34, 3.94) | 0.026 |
| *Race: White* | 0.76 (0.59, 0.98) | 0.125 |
| *Age (in 10 yrs)* | 0.86 (0.77, 0.96) | 0.045 |
| *Compartment (in 10%)* | 1.69 (1.21, 2.37) | 0.026 |
| *HER2/Compartment* | 0.47 (0.19, 1.16) | 0.200 |
| *LumA/Compartment* | 0.66 (0.27, 1.59) | 0.475 |
| *LumB/Compartment* | 0.40 (0.15, 1.02) | 0.146 |
| **Tumor** | | |
| *PAM50: HER2* | 2.51 (1.17, 5.41) | 0.070 |
| *PAM50: LumA* | 0.75 (0.37, 1.50) | 0.450 |
| *PAM50: LumB* | 1.88 (0.90, 3.92) | 0.124 |
| *Race: White* | 0.77 (0.57, 1.04) | 0.124 |
| *Age (in 10 yrs)* | 0.84 (0.74, 0.96) | 0.070 |
| *Compartment (in 10%)* | 1.32 (1.01, 1.74) | 0.101 |
| *HER2/Compartment* | 0.69 (0.48, 1.00) | 0.101 |
| *LumA/Compartment* | 0.87 (0.55, 1.39) | 0.562 |
| *LumB/Compartment* | 0.75 (0.41, 1.38) | 0.446 |
| **Immune** | | |
| *PAM50: HER2* | 1.64 (1.08, 2.50) | 0.096 |
| *PAM50: LumA* | 0.51 (0.33, 0.79) | 0.042 |
| *PAM50: LumB* | 1.30 (0.86, 1.98) | 0.369 |
| *Race: White* | 0.77 (0.59, 1.01) | 0.183 |
| *Age (in 10 yrs)* | 0.84 (0.75, 0.95) | 0.042 |
| *Compartment (in 10%)* | 1.03 (0.70, 1.53) | 0.878 |
| *HER2/Compartment* | 0.48 (0.19, 1.19) | 0.250 |
| *LumA/Compartment* | 1.47 (0.65, 3.33) | 0.494 |
| *LumB/Compartment* | 0.74 (0.29, 1.84) | 0.606 |

Table S3: Hazard ratio estimates, 90% FDR-adjusted confidence intervals, and FDR-adjusted P-values for baseline and compartment-specific interaction Cox models for breast cancer-specific survival.

### 4 Supplemental Figures

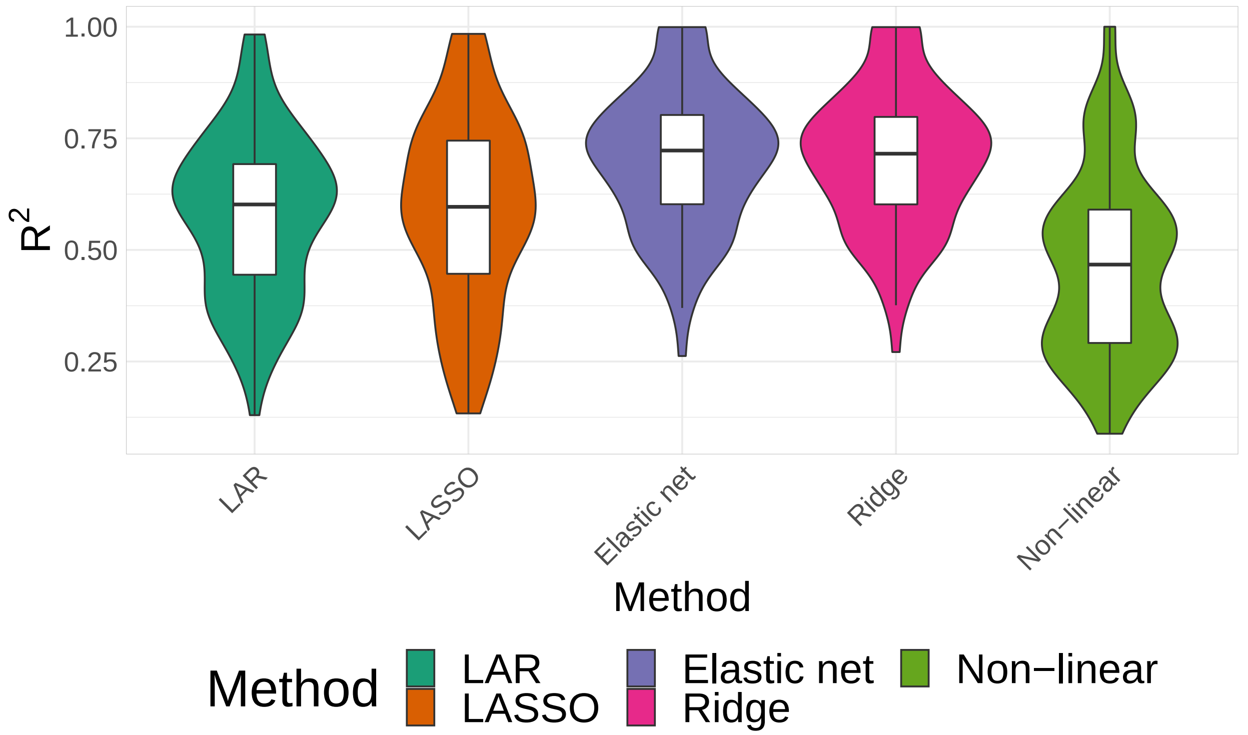


Figure S1: *Comparison of predictive performance of optimization methods used in DeCompress's compressing sensing step.* Violin plots for distributions of cross-validation $R^2$ ($Y$-axis) of the various optimization methods ($X$-axis) employed by DeCompress for compression sensing for 100 randomly selected genes from CBCS. From left to right, least angle regression, LASSO, elastic with $\alpha = 0.5$, ridge regression, and non-linear optimization with $l_1$ norm. Non-linear optimization with either the total variation-adjusted $l_1$ norm or the $l_2$ norm gives similar results as with the $l_1$ norm, and hence is omitted.

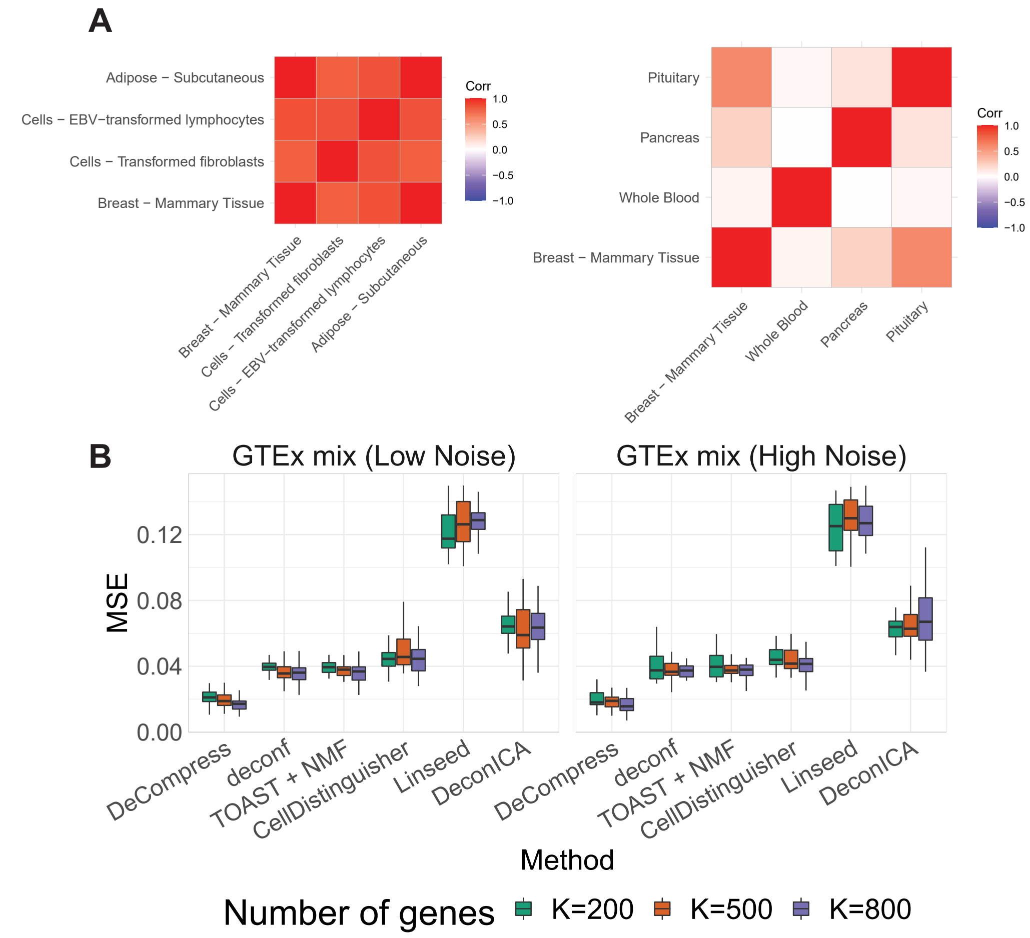


Figure S2: *Impact of varied cell-types on GTEx mixing experiments.* (A) Correlation matrices between tissues used in GTEx *in-silico* matrices. On the left, we show the correlation between tissues used in mixtures in **Figure 2**. On the right, we show the correlation between tissues used in mixtures in **Supplemental Figure S2B**. (B) Boxplots of mean square error ($Y$-axis) between true and estimated cell-type proportions in *in-silico* GTEx mixing experiments across various methods ($X$-axis), with 25 simulated datasets per number of genes. GTEx mixing was done at two levels of multiplicative noise, such that errors were drawn from a Normal distribution with zero mean and standard deviation 4 (left) and 8 (right). Boxplots are colored by the number of genes in each simulated dataset. These simulations were done with 4 cell-types, shown in the right panel of **Supplemental Figure S2A**.

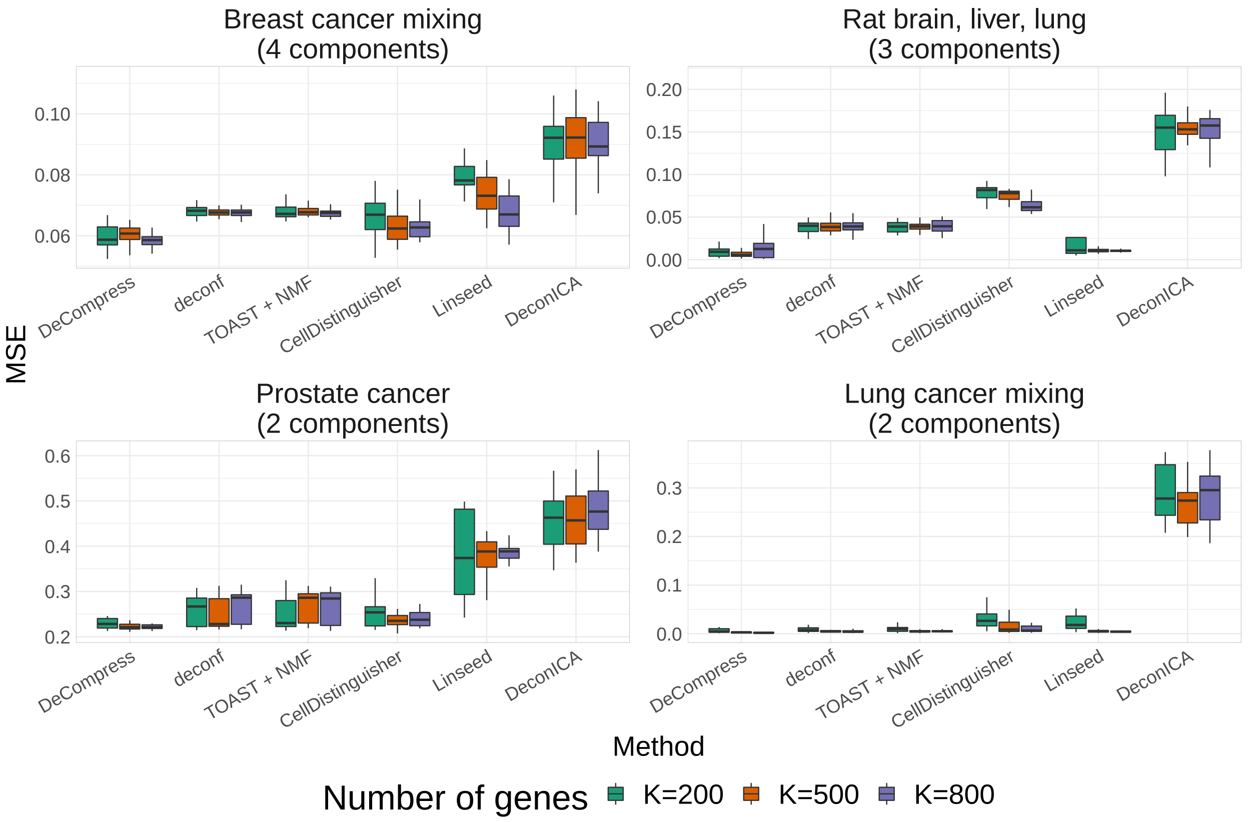


Figure S3: *Benchmarking of deconvolution performance using DeCompress and 5 other reference-free deconvolution in published data examples.* Boxplots of MSE ($Y$-axis) over 25 pseudo-targeted panels using four published datasets over 200, 500, 800, and 1000 genes ($X$-axis). This plot shows the same results as Figure 2C with the addition of DeconICA.

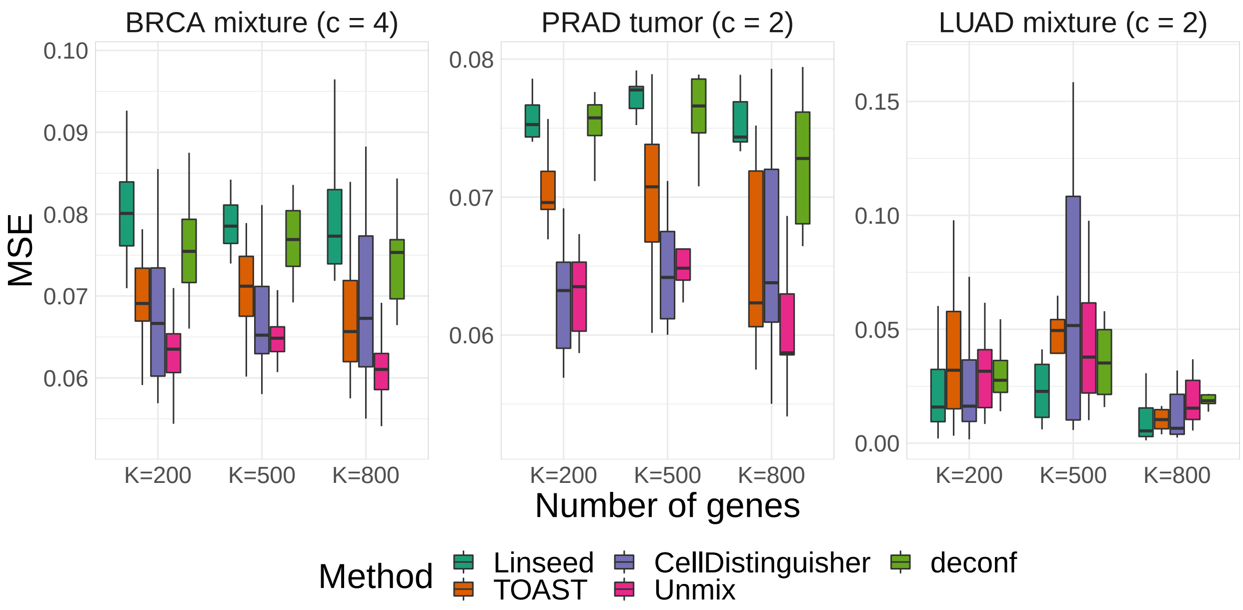


Figure S4: *Comparison of deconvolution performance using decompressed matrix in DeCompress across various methods.* Boxplots of MSE ($Y$-axis) between true and estimated cell-type proportions across pseudo-targeted panels of differing numbers of genes. We compare four reference-free methods (deconf[11], Linseed[1], iterative non-negative matrix factorization with feature selection using TOAST[2], CellDistinguisher[12]) and a reference-based method (unmix[14]) that uses cell-type specific expressions estimated from the reference. Here, we present results from the breast cancer cell line mixtures[24], prostate tumor[25], and lung adenocarcinoma cell line mixtures[26]. We do not include DeconICA[13] in this benchmarking due to large errors across all three datasets.

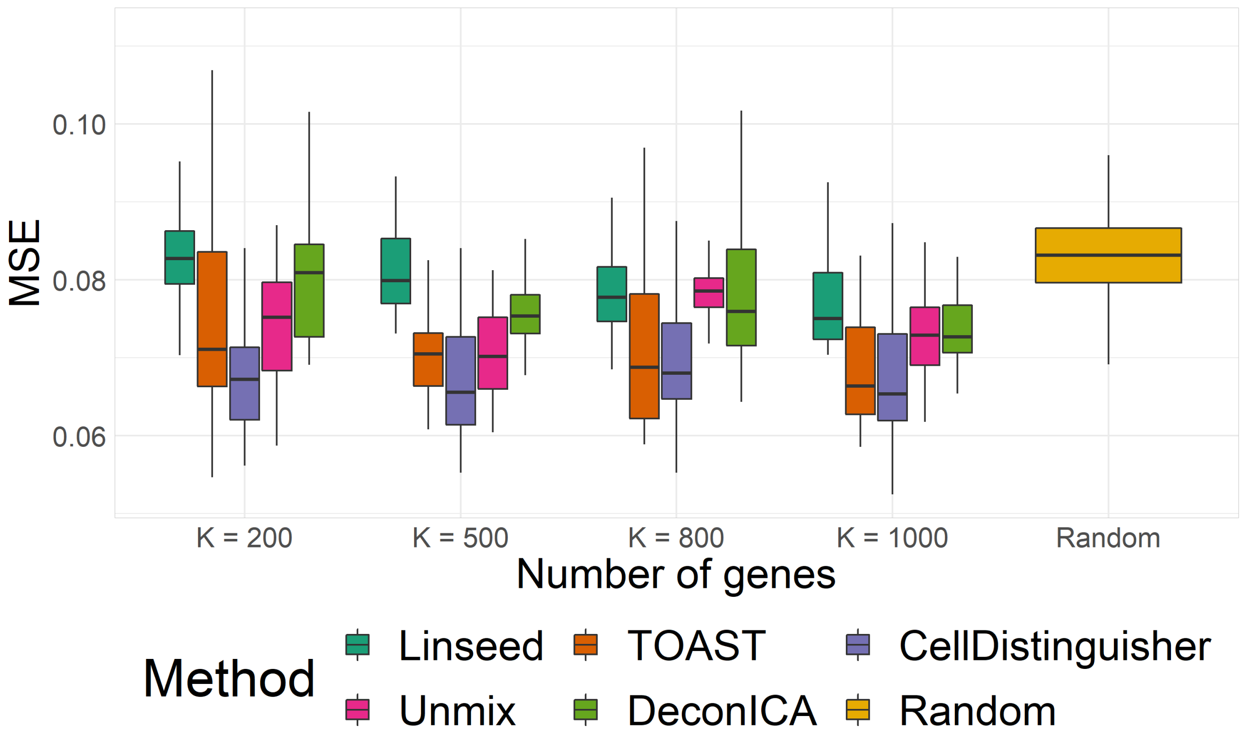


Figure S5: *Deconvolution of breast cancer cell mixture using TCGA-LUAD reference.* MSE ($Y$-axis) across 25 psuedo-targeted panels with different numbers of genes ($X$-axis) of using various reference-free deconvolution methods on decompresed breast cancer cell line data using TCGA-LUAD reference data. The yellow box-plot gives a distribution of the MSE for 1,000 randomly generated cell-type proportions.

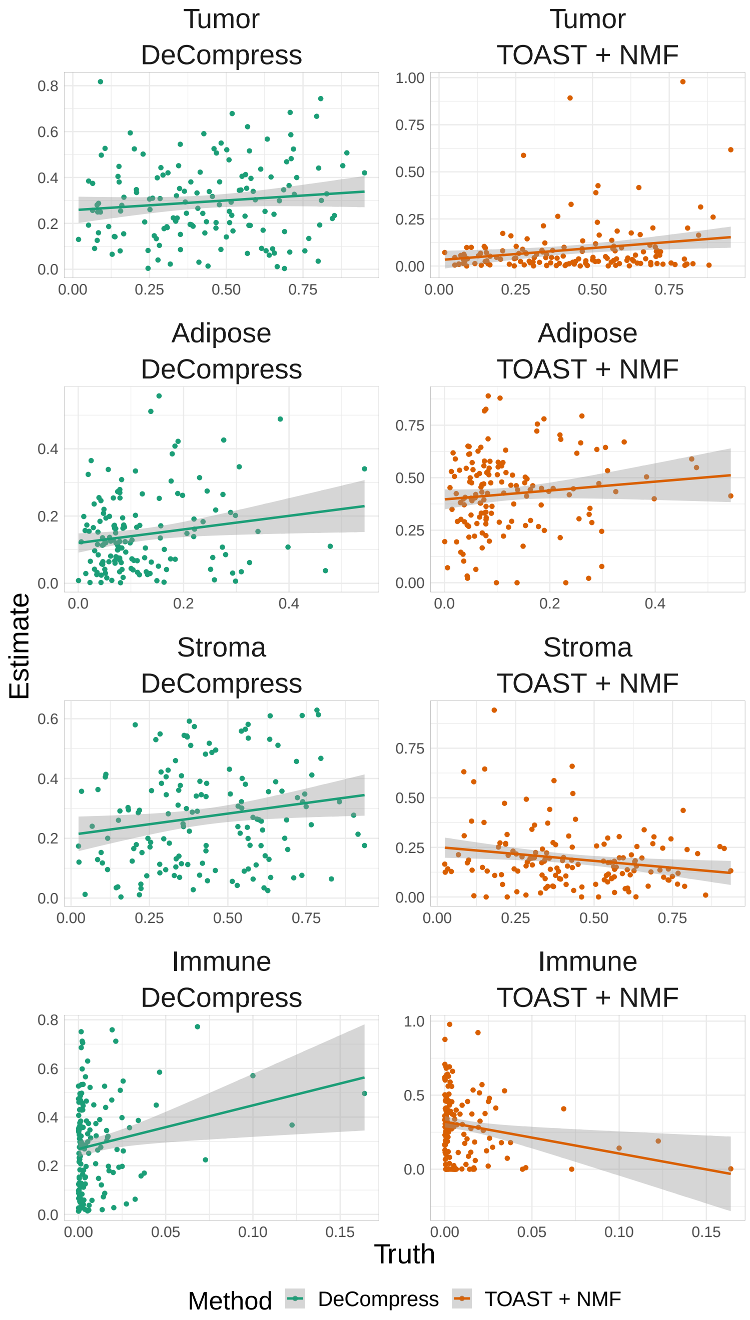


Figure S6: *Scatter-plot of known and estimated cell-type proportions in CBCS using DeCompress and TOAST + NMF.* Plots of true ($X$-axis) and estimated ($Y$-axis) cell-type proportions in CBCS using DeCompress and TOAST + NMF (most accurate benchmarked reference-free method). True cell-type proportions are taken as measurement by a study pathologist for 148 samples. A reference smoothed linear trend line is provided for reference.

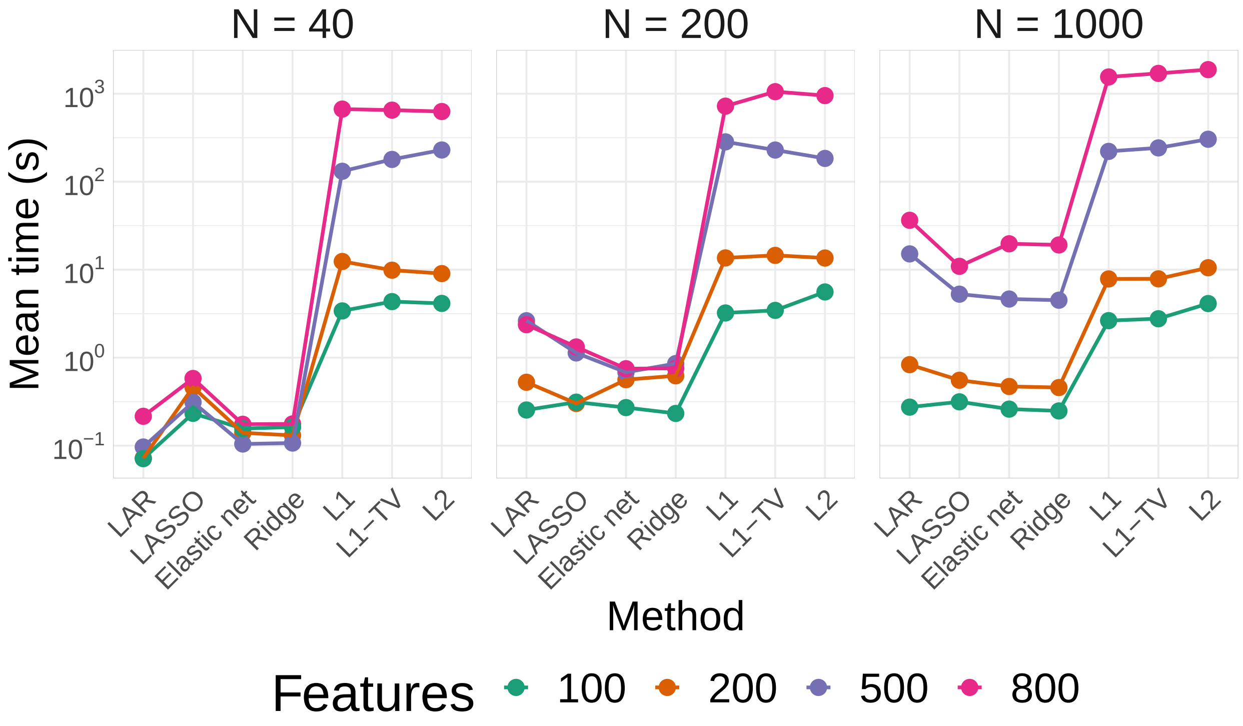


Figure S7: *Comparison of run-times for various methods implemented for compressed sensing in DeCompress.* Over sample sizes of $N = 40$, $N = 200$, and $N = 1000$ and feature sizes of 200, 500, 800, and 100, we plot the mean time of estimation compression model over the 7 methods implemented in DeCompress: least angle regression (LAR), LASSO, elastic net with $\alpha = 0.5$, ridge regression, non-linear optimization with $l_1$ norm, non-linear optimization with total variation-adjusted $l_1$ norm, and non-linear optimization with $l_2$ norm.

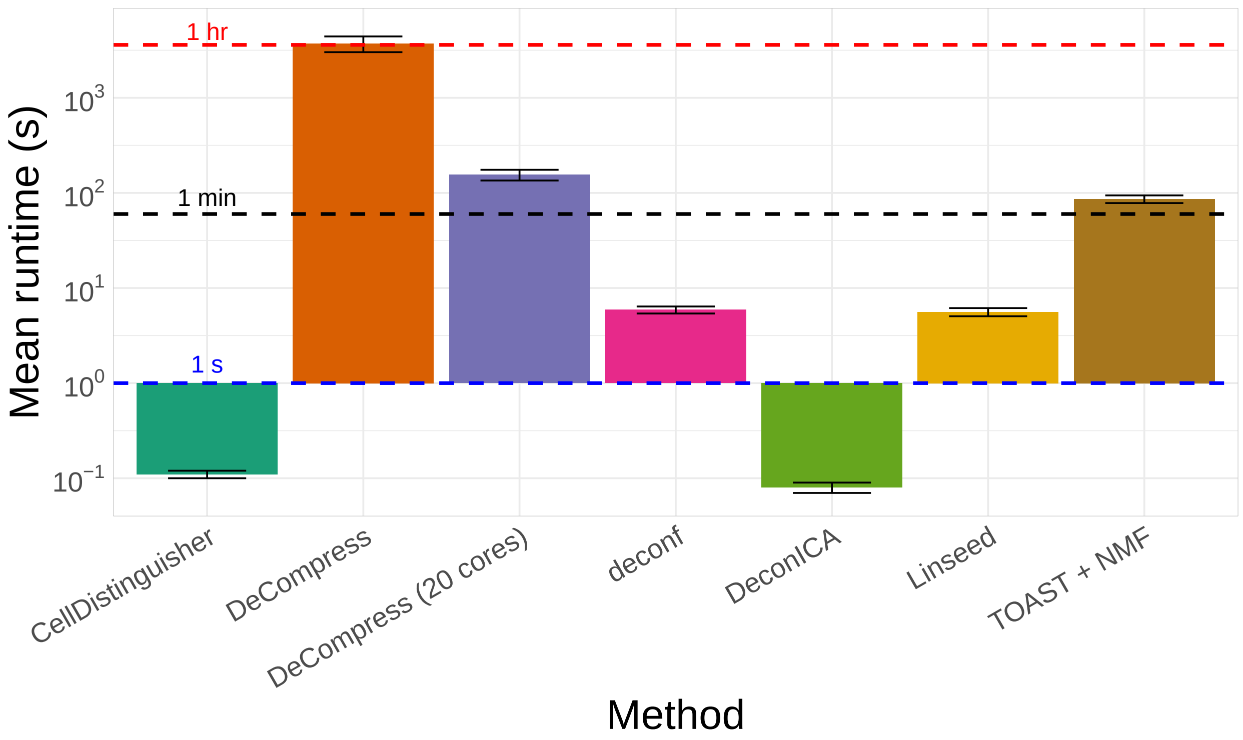


Figure S8: *Comparison of run-times for DeCompress and benchmarked reference-free deconvolution methods.* Mean runtimes in seconds ($X$-axis on logarithmic scale) for methods benchmarked ($Y$-axis): CellDistinguisher, DeCompress (in serial), DeCompress (in parallel with 20 cores), deconf, DeconICA, Linseed, iterative non-negative matrix factorization with feature selection using TOAST. These runtimes were generated by running all methods on CBCS data (1,199 samples with 407 genes). DeCompress was run using TCGA-BRCA (1,212 samples) as a reference. The error bar gives an interval of one standard deviation around the mean runtime. The blue, black, and red dotted lines provide references for 1 second, 1 minute, and 1 hour.

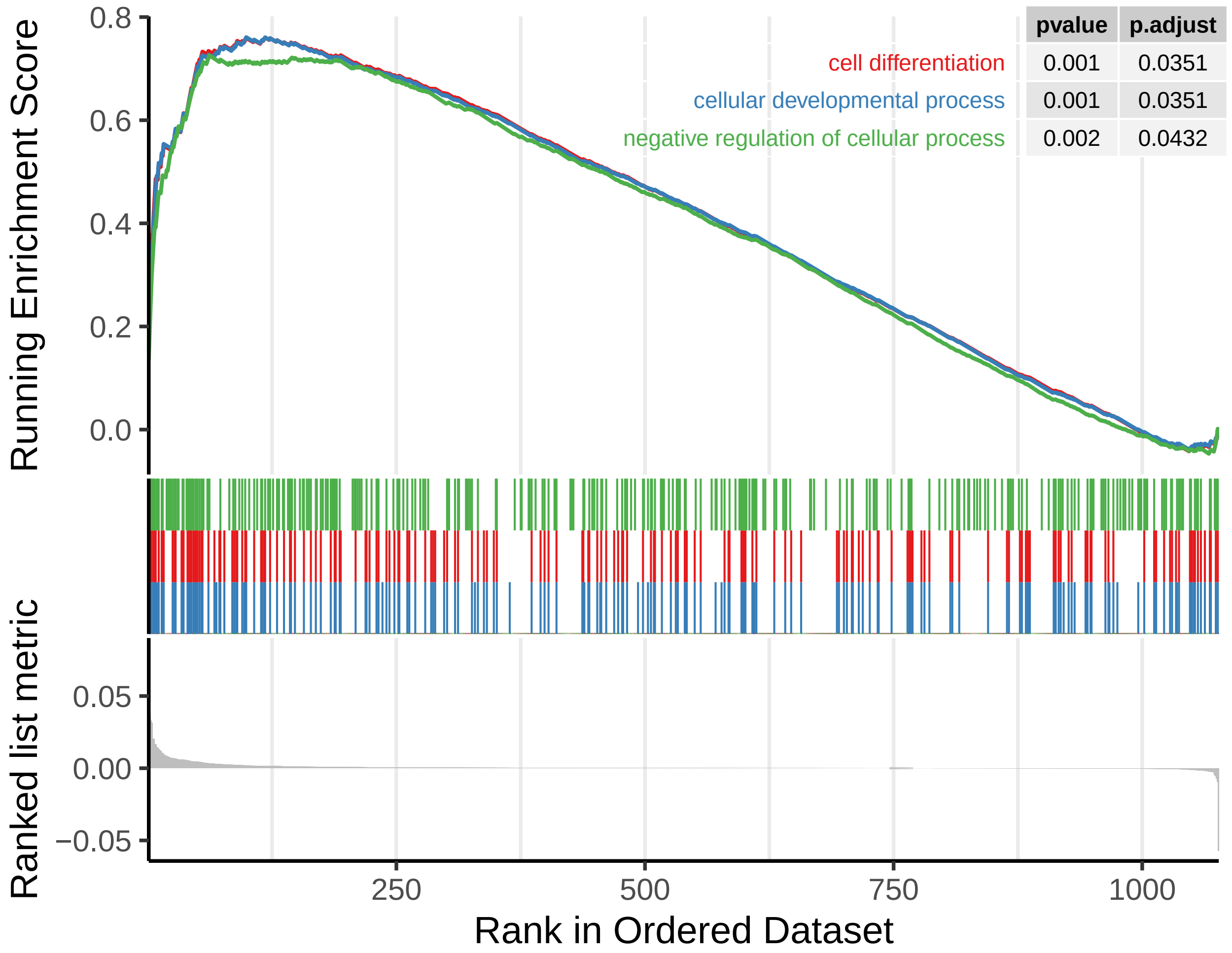


Figure S9: *Gene set enrichment plot for combined C3 and C4 gene signature.* The green, blue, and red lines in the top panel of the plot represents the running enrichment score (ES) for the corresponding gene ontology as the analysis goes down the ranked list. The peak gives the final ES. The green, blue, and red lines in the middle of the plot shows where the members of ontological groups in the dataset first appear in the ranked list. The bottom panel shows the value of the ranking metric as it moves down the list of the ranked genes.

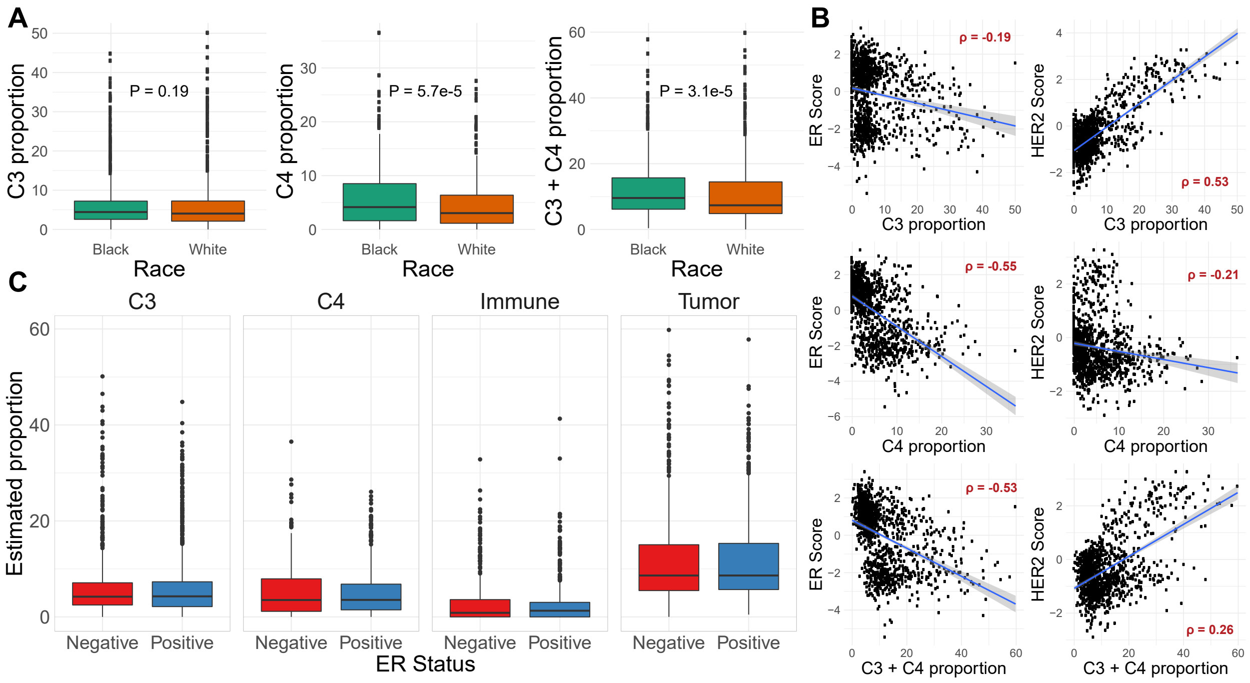


Figure S10: *Comparison of compartment proportion estimates with race and different clinical subtype metrics.* (A) Boxplot of C3, C4, and C3 + C4 proportions across race with $P$-value of Wilcoxon rank-sum test provided. (B) Scatterplot of compartment proportions ($X$-axis) and ER or HER2 score from PAM50 classification algorithm. A regression line is provided with a Spearman correlation $\rho$ for reference. (C) Boxplot of C3, C4, immune, and tumor compartment estimates acros clinical ER status.

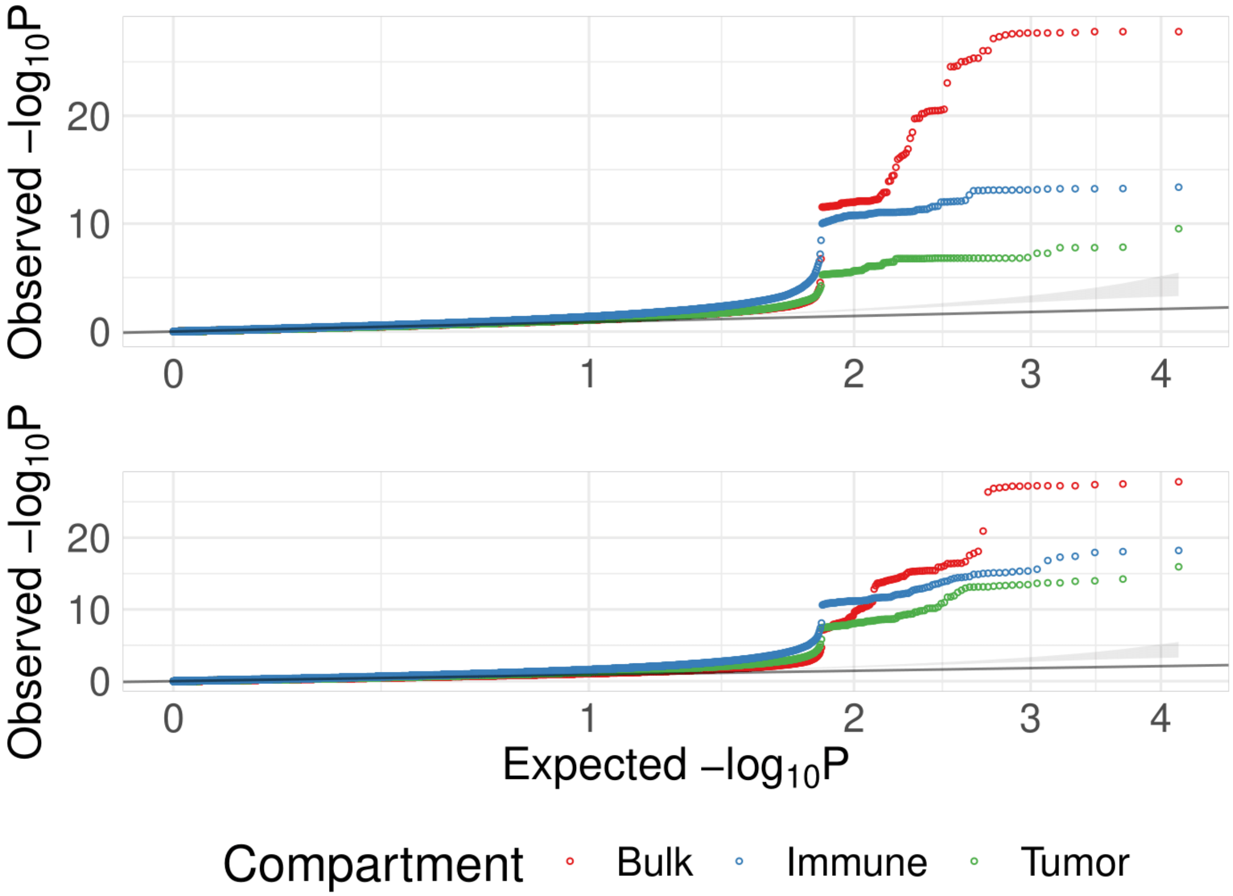


Figure S11: *Manhattan plot of cis-eQTLs across the genome in AA CBCS samples.* $-log_{10}P$-values of eQTL association ($Y$-axis) across chromosomal position of *cis*-eQTLs across bulk (top), immune (middle), and tumor (bottom) models. Top *cis*-eGenes are labelled.

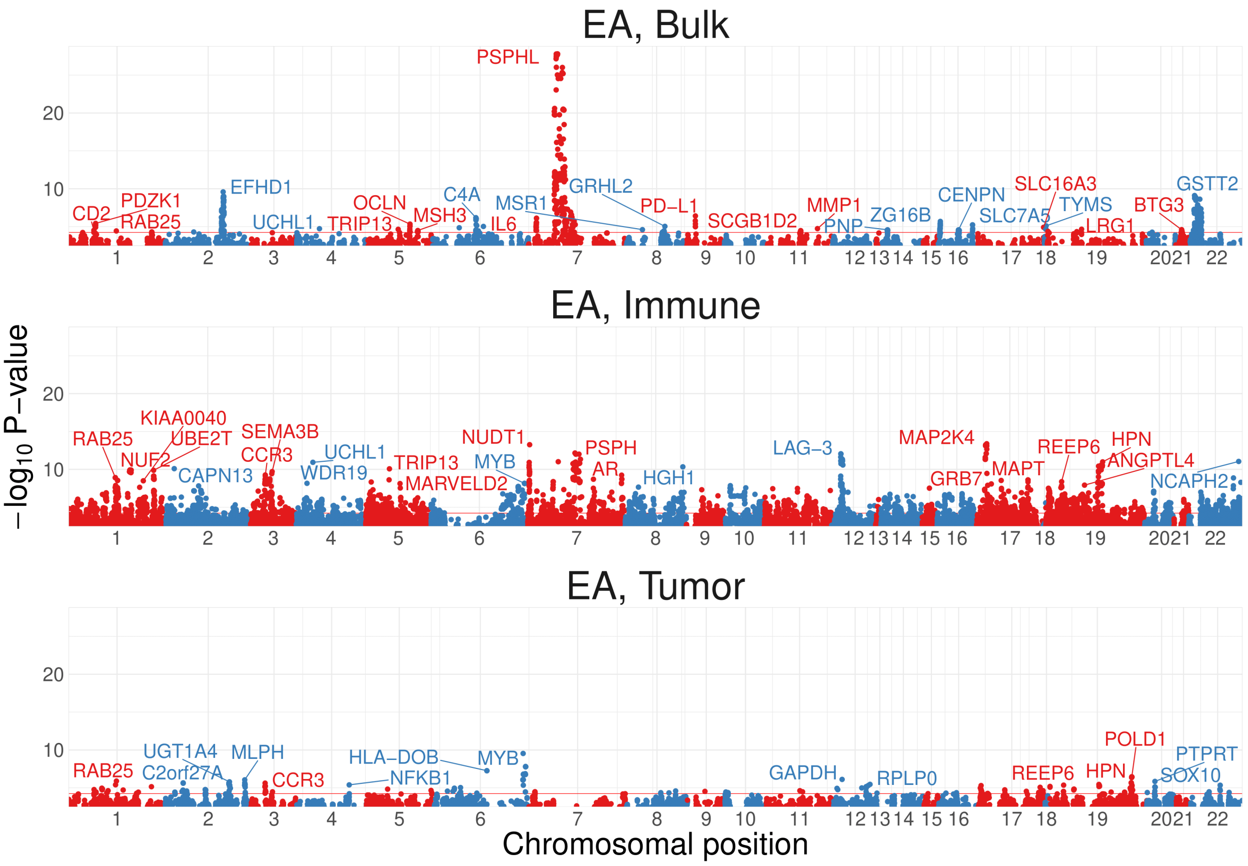


Figure S12: *Manhattan plot of cis-eQTLs across the genome in EA CBCS samples.* $-log_{10}P$-values of eQTL association ($Y$-axis) across chromosomal position of *cis*-eQTLs across bulk (top), immune (middle), and tumor (bottom) models. Top *cis*-eGenes are labelled.

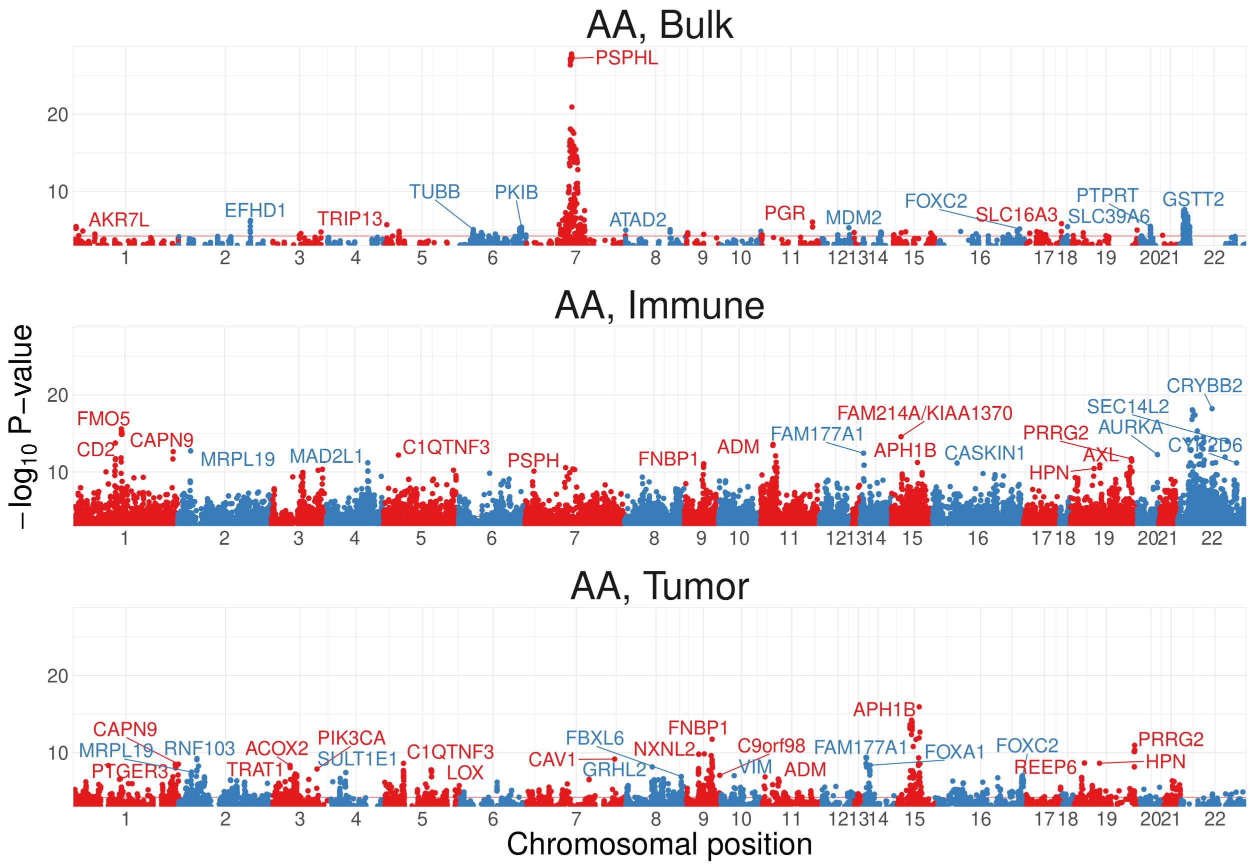


Figure S13: *Manhattan plot of cis-eQTLs across the genome in AA CBCS samples.* $-log_{10}P$-values of eQTL association ($Y$-axis) across chromosomal position of *cis*-eQTLs across bulk (top), immune (middle), and tumor (bottom) models. Top *cis*-eGenes are labelled.

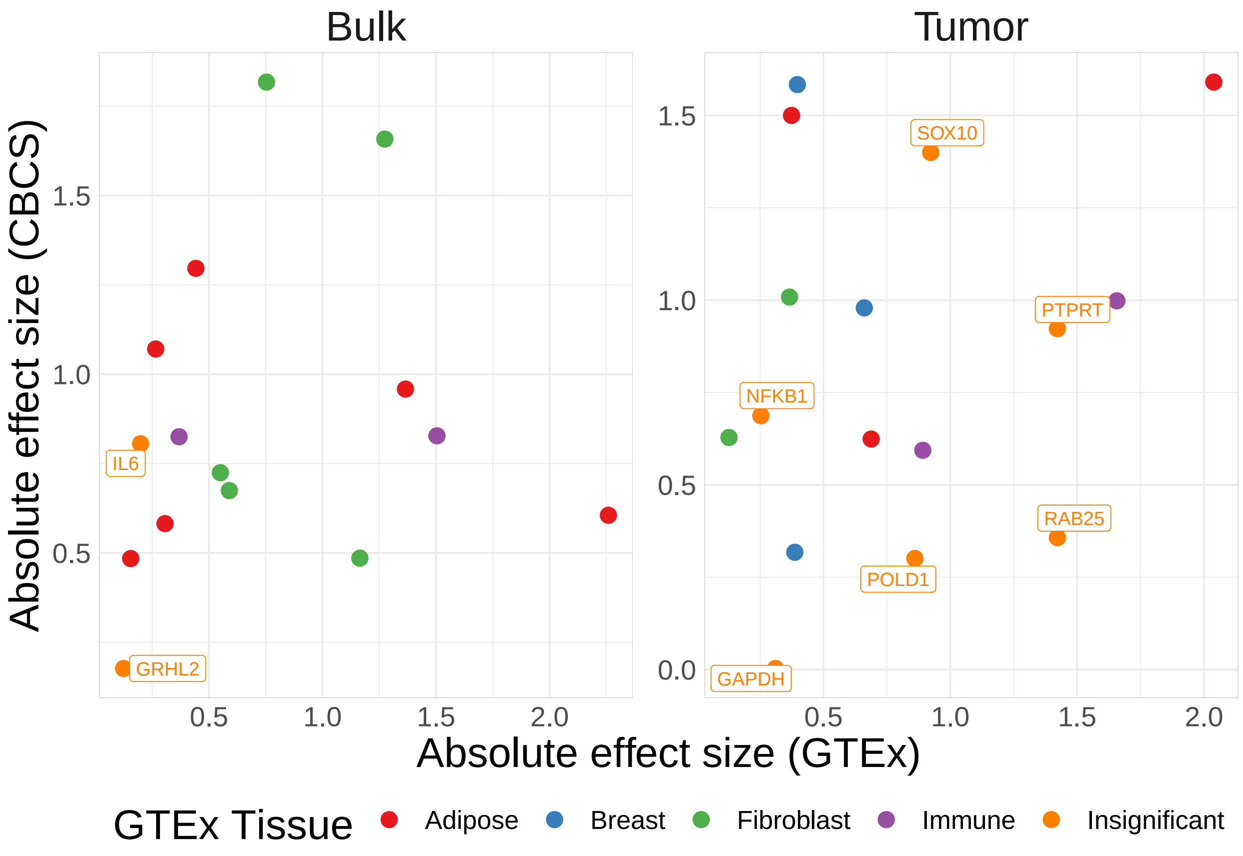


Figure S14: *Cross-referencing of bulk and tumor-specific CBCS EA cis-eGenes with GTEx.* Comparison of absolute effect sizes of eGenes with significant *cis*-eQTLs in EA CBCS ($Y$-axis) and GTEx ($X$-axis) over tissue type, stratified by bulk and tumor-specific eQTLs. eGenes are colored by the GTEx tissue that shows the eQTL with smallest $P$-value.

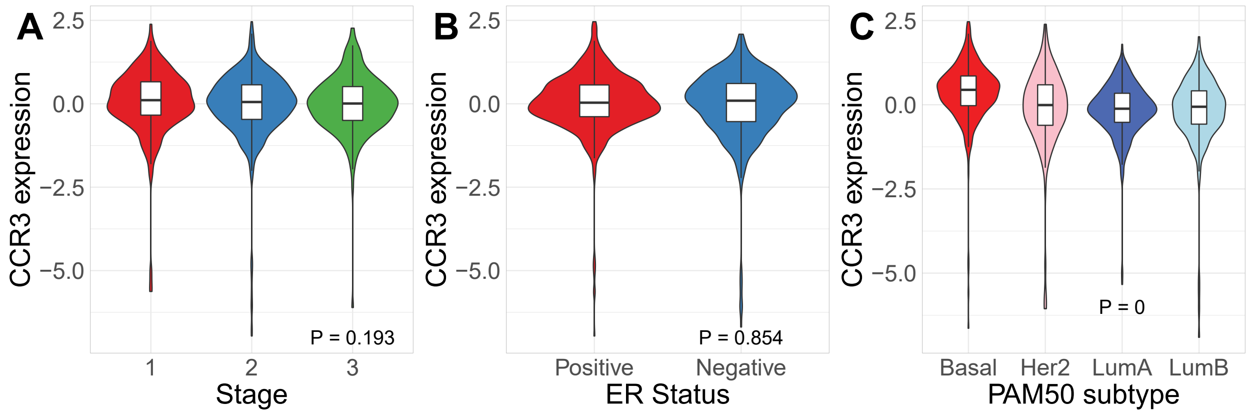


Figure S15: *Associations of CCR3 expression across clinical variables, subtypes, and mortality.* Violin plots of *CCR3* expression across breast tumor stage (A), estrogen status (B), and PAM50 molecular subtype (C).
